# HUGMi: Human Uro-Genital Microbiome database and hybrid classifier for improved species level annotation of 16S rRNA amplicon sequences

**DOI:** 10.1101/2025.05.01.651608

**Authors:** Debaleena Bhowmik, Sandip Paul

**Author notes:** To whom correspondence should be addressed: Sandip Paul, JIS Institute of Advanced Studies and Research Kolkata, JIS University JIS School of Medical Science & Research Campus, 51, South Nayabaz, GIP Colony, Santragachi, Howrah 711112, West Bengal, INDIA.

## Abstract

The human urogenital microbiota encompasses commensal microorganisms harbouring the urinary and reproductive tracts, and has been associated with the pathophysiology of multiple urogenital disorders. Here, the Human Urogenital Microbiome (HUGMi) workflow is introduced, wherein a manually curated, niche-specific, 16S rRNA database is integrated with a novel hybrid classification algorithm to enhance amplicon-based species-level taxonomic resolution in urogenital microbiome investigations.

The HUGMi database is curated through rigorous quality assessment of sequences, nomenclature correction of taxonomies and meticulous selection of bacterial species biologically significant to the human urogenital region. Quantitative evaluation of taxonomic assignment with mock communities demonstrates that HUGMi consistently achieved superior accuracy compared to conventional databases, with statistically significant improvement in F1 scores. Comparative analyses across three sets of biological data also reveal a robust and versatile performance, with both QIIME2 BLAST and sklearn-based taxonomic classifiers and with varied 16S regions. Following these outcomes the HUGMi hybrid classifier was developed that combines the specificity of BLAST-based methods with the probabilistic framework of learning-based classifiers. This approach enabled superior species-level identification through varying confidence thresholds. In contrast to the limited number of species identified in real-time studies, the integrated outcome of the HUGMi database and hybrid classifier, not just replicated the original results but simultaneously revealed a spectrum of previously unidentified species.

Despite its crucial role in human health, the urogenital microbiome is comparatively understudied with respect to other body sites. The HUGMi pipeline addresses this empirical lacuna by identifying a substantially expanded range of species-level taxa. This enhanced resolution will not only augment downstream functional studies, but also aid in the investigation of specific bacterial influence on urogenital homeostasis and disease pathogenesis.

## INTRODUCTION

Microbial communities are integral to a wide range of biochemical processes and ecological interactions, making them crucial for both human and environmental health. Historically, microbiome research has solely focused on identifying the presence-absence of microbes, and from there estimated the metabolic functions, niche specific roles, or correlation with various ecological or clinical states. Although an overview of the nature and roles of microbes could be predicted with the genus level taxonomic recognition, the accurate ecological function and metabolic capabilities can only be derived through species level identification (Martiny et al., 2015).

Metagenomic analysis predominantly relies on 16S amplicon sequencing, which is an economically viable technology for yielding high-throughput data. Amplicon data is subsequently grouped into Operational Taxonomic Units (OTUs) or Amplicon Sequence Variants (ASVs), through clustering methods like USEARCH (Edgar, 2010) or denoising algorithms like Deblur (Amir et al., 2017), DADA2 (Callahan et al., 2016), or UNOISE3 (Edgar, 2016). These OTUs/ASVs are then assigned with taxonomies by employing a reference database and taxonomic classifiers like RDP classifier (Cole et al., 2014) and classifiers present in QIIME2 (Bolyen et al., 2019). Over the years the 16S reference databases like Genome Taxonomy Database (GTDB) (Parks et al., 2022), NCBI RefSeq 16S database (O’Leary et al., 2016), Greengenes (DeSantis et al., 2006; McDonald et al., 2024) and SILVA (Quast et al., 2013) have undergone extensive upgradation in order to include more sequences and improve species-level clarity. However, they still exhibit some limitations, such as inconsistent nomenclature, artifacts, and uncertain taxonomic assignments at various levels (species, genus, family, order, and class).

Alongside these comprehensive databases, there has been development of niche-specific databases as well, including HITdb (Ritari et al., 2015), eHOMD (Chen et al., 2010), RIM-DB (Seedorf et al., 2014), DictDb (Mikaelyan et al., 2015), MiDAS (McIlroy et al., 2015), AutoTax (Dueholm et al., 2020), and TaxAss (Rohwer et al., 2018). The key advantages of these ecosystem-specific databases are enhanced sensitivity to the microbes specific to the ecosystem, decreased false positives and a monophyletic reference structure. The use of these specialized databases leads to more precise taxonomic assignments compared to the broader, general-purpose databases mentioned earlier (Rohwer et al., 2018).

Despite the proliferation of niche-specific amplicon databases, they are generally directed towards environmental ecosystems or human gut, barring the one for the human oral cavity. Few databases have focused on the female vaginal region (Holm et al., 2024; Molano et al., 2024), however there is a significant gap in repositories for the entire human urogenital microbiota (UGM). The UGM refers to the community of microbes that naturally reside in the human urinary and genital tracts, including the urinary, periurethral, vaginal, and penile microbiomes, and the idea of a native microbial community in the urinary system is relatively recent. It is found to play a key role in modulating susceptibility towards certain uropathogens and the severity of infection caused in urinary tract infections (UTIs) and urethritis (Perez-Carrasco et al., 2021); they have also been associated with interstitial cystitis (Xu et al., 2021), bladder pain syndrome (IC/BPS) (Bhide et al., 2020), bladder cancer (Bučević Popović et al., 2018) and prostate cancer (Pernigoni et al., 2023). In contrast, the vaginal microbiome is a subcategory of the UGM that has been extensively studied. The five community state types (CSTs) of the vaginal microbiome are well established (Bedford et al., 2020), however there has been reported connection with the urogenital microbiome also (Thomas-White et al., 2018). This highlights the importance of bridging the knowledge gap regarding the UGM, which lead to the development of the Human Uro-Genital Microbiome (HUGMi) workflow: a 16S database for increased taxonomic precision in urogenital datasets and a hybrid classifier for improved taxonomic annotations.

The HUGMi database is a secondary, manually-curated, niche-specific database, with sequences and taxonomies extracted from Genome Taxonomy Database (GTDB) (Parks et al., 2022), NCBI RefSeq 16S database (O’Leary et al., 2016), Greengenes2 (GG2) (McDonald et al., 2024), SILVA (Quast et al., 2013), Ribosomal Database Project (RDP) (Cole et al., 2014) and EzBioCloud 16S database (Yoon et al., 2017). It is envisioned that the HUGMi framework aids in urogenital microbiome research by precise bacterial identification and eventual functional delineation. And this might further advance the development of new therapies targeting the urogenital microbiota, their community structure and role in pathogenesis of urologic diseases.

## MATERIALS AND METHODS

The diagrammatic representation of the methodology for creating of the HUGMi database is demonstrated in Figure 1. The step-by-step materials and methods are detailed below.

**Figure 1:**
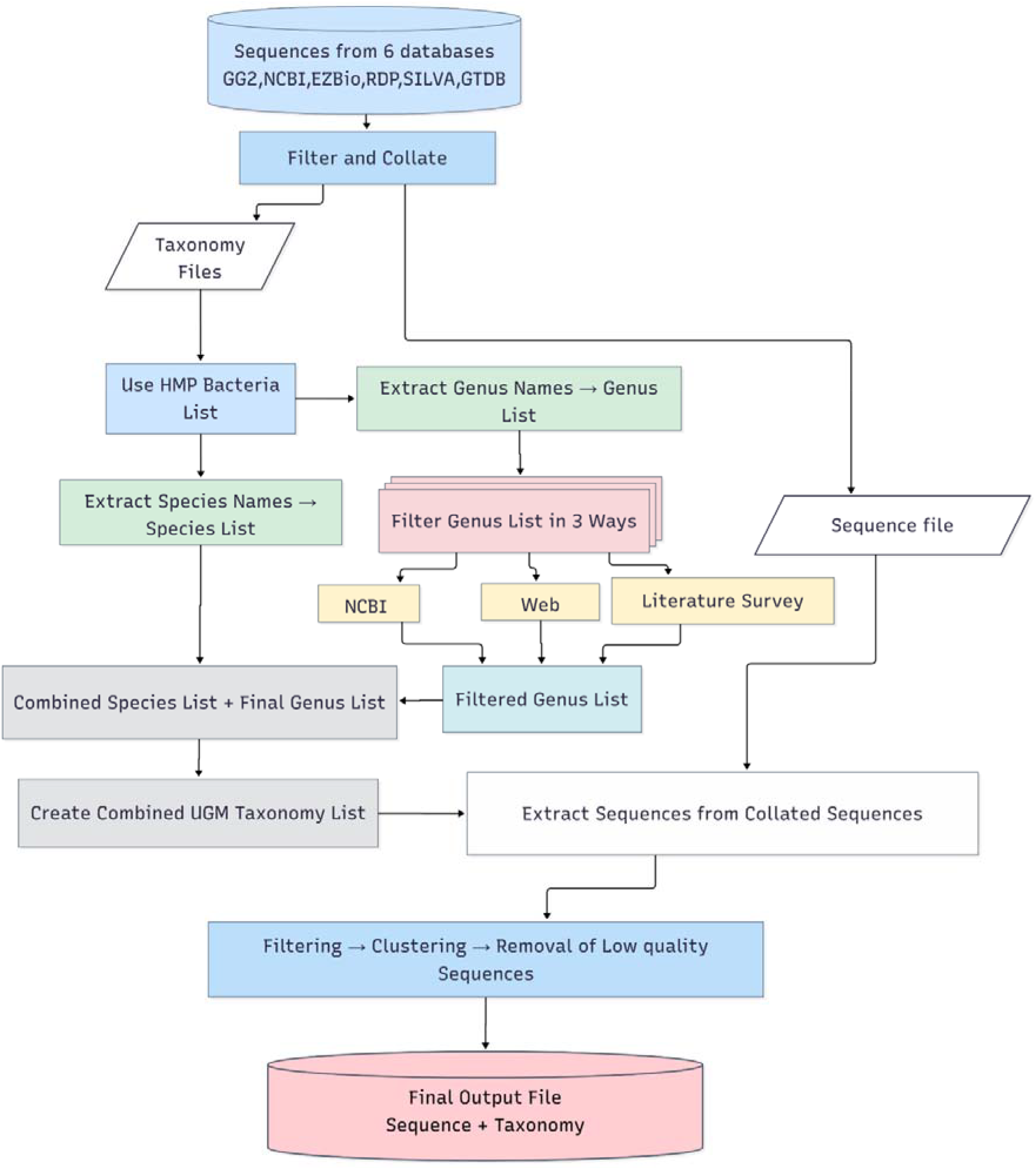
Overall flowchart depicting the methodology for curating the HUGMi database.

**Figure 2:**
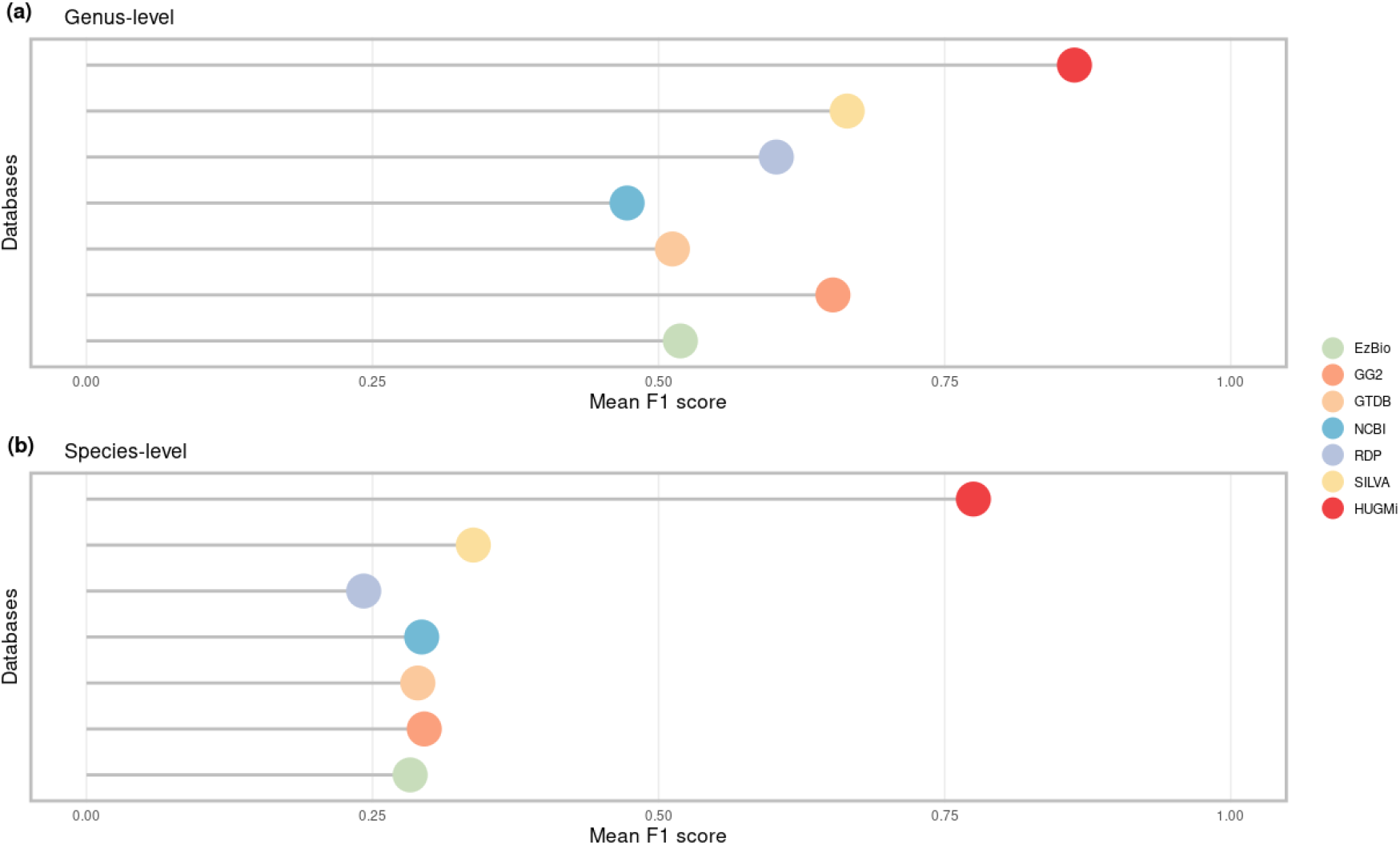
Lollipop plot illustrating the mean F1 score across databases for mock data.

### Curation of the HUGMi database

#### I. Cumulation of 16S rDNA sequences and taxonomy

The HUGMi database essentially is a manually curated, cross-referenced, secondary database sourced from six of the popular microbial databases for 16S amplicon sequence analyses - Genome Taxonomy Database (GTDB) (v220), NCBI RefSeq 16S database, Greengenes2 (GG2) (v2022.10), SILVA (v138), Ribosomal Database Project (RDP) and EzBioCloud 16S database. The primary aim for this database was to acquire species-level identification of bacteria and in order to achieve that goal, these databases were filtered with exclusion criteria of archaeal, incomplete (not having assigned names up to species-level), unclassified or putatively named sequences. There are a significant percentage of putative nomenclatures in the GTDB, Greengenes2, SILVA, EzBioCloud and RDP databases, which were consciously excluded as they would not have aided in the bacterial identification. Once all these criteria were fulfilled, the extracted sequences, totaling up to 810989 were collated into a single fasta file along with a collated (.tsv) file for associated taxonomies. The number of taxa of the constituent databases has been detailed in Supplementary Table 1.

#### II. HMP urogenital microbial taxonomy extraction

The Human Microbiome Project (Turnbaugh et al., 2007) harbors a plethora of data concerning the human microbiome including the different populace of bacteria residing in different body sites. From the *HMP1 Metadata Project Catalog* webpage, a comprehensive list of all bacterial genera and species, specific to the human urogenital region was obtained and it encompassed all the related body sites for both male and female. In order to create a niche specific urogenital database, this particular set of 492 bacterial names served as reference points of sequence and taxonomy inclusion from the collated file created in the previous step (I.). Although the set contained many names of bacteria with “Genus-species designations”, there were quite a few of them with only Genus-level identifications. In the former case, all related taxonomies were extracted from the collated taxonomy file; while in case of the latter, all possible species belonging to those genera were extracted from the same file, and then subjected to another level of filtration downstream.

#### III. Targeted selection of niche specific microbial taxonomies

The “Genus-only” list of urogenital bacteria obtained from the HMP website, was used to extract all available species taxonomy associated with those genera, however not all the species from those genera are niche specific (found in human urogenital region) or biologically relevant (associated with the urogenital region in some biological form, like disease states). In order to identify only the niche specific, biologically relevant species, a three-pronged approach was developed (Figure 1), and is explained as follows:

##### a) Automated extraction of taxonomic ecological data from web-resources

This part of the search was implemented through a Python-based web scraping program using Selenium (v4.24) and chromedriver (version 127.0.6533.72). Employing predefined taxonomic search terms, the program systematically queried web-resources and extracted specific HTML elements identified by their respective class attributes. These HTML elements contained ecological data relevant to a particular bacterial name, and were stored in a tabular format. From there, the bacterial species specific to the urogenital region were isolated manually.

##### b) Systematic retrieval of biological source using NCBI’s Entrez Programming Utilities (E-utilities)

The esearch and efetch programs of the NCBI E-utilities (eutils) (Sayers, 2022) aided in gathering the isolation sources of the concerned bacteria, from the NCBI repository. With the help of xml.etree.ElementTree module, a Python program first accessed the esearch program with parameters {db=nucleotide, term=*input_microbe_name*, retmode=json} for automated search; next using the efetch program with parameters {db=nucleotide, id=ids, rettype=gb and retmode=xml} detailed records for bacterial species were fetched. The program then parsed through each of the resulting XML files selecting the element that contained the isolation source {GBQualifier_name=isolation_source}, yielding a tabulated output. The bacterial species linked with the urogenital niche were manually separated.

##### c) Computational literature mining

The PubMed archives of NCBI were mined to collect research articles focusing on urogenital microbiome and diseases, with the help of MeSH (Medical Subject Headings) search terms, as described in Supplementary Table 2. The URLs of the papers were collected and once again a Python-based web scraping program using BeautifulSoup (v4.12.3) was employed for parsing the HTML webpage and extracting names of bacterial species relevant to the human urogenital body site.

The computational assortment of relevant bacteria was supplemented by manual bibliographic extraction resulting in a concatenated list of bacterial names (both genus and species names). This, along with the HMP “Genus-species designated” list of bacterial names obtained in step (II.), constituted a complete repertoire of bacterial species of the human urogenital microbiome. This repertoire contained 1,033 unique taxa, and was eventually used to extract the sequences and taxonomies from the collated files created in step (I.). The resultant files constituted 3,58,677 sequences and taxa.

#### IV. Quality assessment of the extracted sequences and nomenclature correction of the taxonomies

The quality of extracted sequences in step (III.), were assessed using two factors: the sequence length distribution and ambiguous base frequency. With the help of two separate Python programs the extracted sequences were filtered, with inclusion criteria of sequence length between 900-1700 nucleotides, and zero ambiguous base (only sequences with ATGC base content included). This yielded 2,94,711 sequences.

The changes in bacterial nomenclature made by the ICNP (International Code of Nomenclature of Prokaryotes) as well as via recent publications were incorporated in the 2,94,711 taxonomies, to ensure consistency and non-redundancy. In order to establish that, a Python program using BeautifulSoup (v4.12.3) was developed, which first performed automated web-search of bacterial names in the NCBI Taxonomy Browser, and parsed through the output to extract the updated nomenclature from NCBI for each of the taxa. Changes were incorporated throughout the seven taxonomic levels and not just at the Genus-species level.

#### V. Unsupervised spurious sequence detection for the uniformity of sequences and respective taxonomies

The final sanity check for the HUGMi database was to ensure that the member sequences are coupled with the correct taxonomy. Since it was a secondary database compiled from other databases, there was an absence of reference sequence. Thus, an unsupervised method, which did not require any reference point, was developed to remove mismatched sequences with the taxonomies. The first step included sequence clustering of the entire database at 99% identity and 99% coverage, using MMseqs2 (Steinegger & Söding, 2017). Then with the help of the Biopython package and Python programming, clusters showing mismatch between representative taxa and member taxa were eliminated. After the removal of the mismatched sequences and taxa, there was a loss of 53 unique taxa from the urogenital repertoire. To partly salvage this loss, sequences with adequate quality and belonging to the missing taxa were manually added. This finally yielded 1,017 unique taxa and 1,92,254 sequences.

### Datasets for the validation of HUGMi database

#### Designing randomized mock data for accuracy calculation

In order to verify the performance of the HUGMi database, 5 mock datasets specific to the urogenital region were created with the help of a Python program. The program first randomly selected 50 taxa from the 1017 names of urogenital bacteria repertoire and then extracted one random sequence each from the collated sequence file (step (I.)) for each of the taxa, ensuring that the selected sequence does not contain any ambiguous base. For negative control, a fasta file of 5 random sequences belonging to no organism (in parts or whole) was created, and merged with each of the datasets, resulting in 5 mock datasets of 55 sequences each. Taxonomic assignment for all the mock datasets were performed using HUGMi as well as all the constituent databases. The F1 score was then calculated as per Supplementary Figures 1 and 2.

#### Selection and processing of publicly available datasets

To evaluate the taxonomic assignment capability of the HUGMi database with real time data, three studies with publicly available amplicon sequences were selected. Dataset I was a study by Popović et al. (2017) (Bučević Popović et al., 2018), who studied the urine microbiome from urine samples of patients with Bladder cancer and healthy individuals. For dataset II, research by Ilhan et al. (2019) (Ilhan et al., 2019) was selected and they studied the vaginal microbiome of cervical cancer patients as well as individuals with cervical dysplasia and HPV infection. Due to the limited number of report taxa in this case, the raw files were separately processed with USEARCH (Edgar, 2010). Closed-reference OTU picking was implemented with Greengenes (v13.8) (DeSantis et al., 2006) database aligning with the original methodology. The final dataset (III) was from Musa et al. (2023) (Musa et al., 2023) study of vaginal microbiome from cervicovaginal lavage of patients with same medical dispositions as of dataset II. Supplementary Table 3 provides additional information regarding the datasets, including their accession numbers. All three datasets were processed using standard QIIME 2 denoising protocol using the dada2 plugin.

For datasets I and II differential analysis was performed after taxonomic assignment with HUGMi, for extracting significant taxa belonging to each of the groups in the respective studies. Phyloseq (McMurdie & Holmes, 2013) and DESeq2 (Love et al., 2014) R packages were used for this analysis.

### Methods of taxonomic annotation using HUGMi database

#### Employing qiime2 feature-classifier for taxonomic assignment

Taxonomic assignment of the mock and validation datasets was implemented using two of the qiime2 (v2024.10) feature-classifers, i) classify-consensus-blast and ii) classify-sklearn. In case of the BLAST method, first a BLAST database (BLASTDB) was created from the concerned database using the q2 makeblastdb plugin and then the classify-consensus-blast with parameters {perc-identity=1.0, query-cov=0.95, maxaccepts=10} was used for taxonomy assignment. In case of the sklearn based approach, first the databases were modified (quality control and 16S region specific trimming) using q2 RESCRIPt plugin (Ii et al., 2021) and then trained a naive bayes classifier using default parameters. Finally with the help of the q2 classify-sklearn and the classifier, taxonomy was assigned. In this case three permutations of the confidence parameter were used {0.7(default), 0.8 and 0.9}.

#### Development of a qiime2-specific hybrid classifier

To extract the highest number of taxonomic assignments as well as maximize on the specificity for a given amplicon sequencing data, a hybrid approach was formulated. This hybrid classifier is a Python-based program that is compatible with qiime2 (q2 version >2023.7) and it assigns taxonomy using both BLAST as well as sklearn methods. It amalgamates the specificity of BLAST with the robustness of learning-based classifiers, achieving not only accuracy but also an increased number of taxonomic assignments, then either of the methods alone. The program performs the BLAST classification first, and then extracts all the ASVs which either remained unassigned, or received a taxonomic assignment lower than the genus-level (e.g. family, class). With this set of ASVs, the sklearn classifier is employed. The parameters are all user-dependent giving them the freedom to modulate the stringency of this classifier based upon their experimental requirements.

## RESULTS

### Benchmarking the precision of taxonomic assignment using defined synthetic communities

To quantitatively evaluate HUGMi’s annotation precision, relative to the other databases, the F1 score metric was employed. The True Positive (TP), True Negative (TN), False Positive (FP), and False Negative (FN) values for taxonomic classification were calculated according to the confusion matrix in Supplementary Figure 1 and subsequently, the Precision, Recall and F1 scores were calculated using the equations in Supplementary Figure 2. These scores were calculated for all the databases on all 5 mock communities. The average F1 score for the 5 mock communities were calculated for each of the databases and the results are depicted in Figure 2. The F1 scores of the HUGMi database were significantly higher than the remaining, both at the genus and species-level annotations. Pairwise t-test affirms the same outcome (Supplementary Table 4).

### Assessing the efficacy of HUGMi database using clinical microbiome samples

#### Taxonomic annotation of dataset I using QIIME2 BLAST classifier

The fundamental aim of developing the HUGMi database was to enhance taxonomic resolution, especially at the species level, from amplicon sequencing itself. The implementation of QIIME2 (Bolyen et al., 2019) consensus blast classifier (Camacho et al., 2009) contributes to this objective by offering higher specificity in taxonomic assignment. This increased differential ability enables precise identification of taxa, thus improving the overall accuracy and reliability of the taxonomic classifications. BLAST classification results in Figure 3 show comparable taxonomic assignment counts at both phylum and genus level with respect to other databases; however at the species level HUGMi outperforms all of them (Supplementary Table 5).

**Figure 3:**
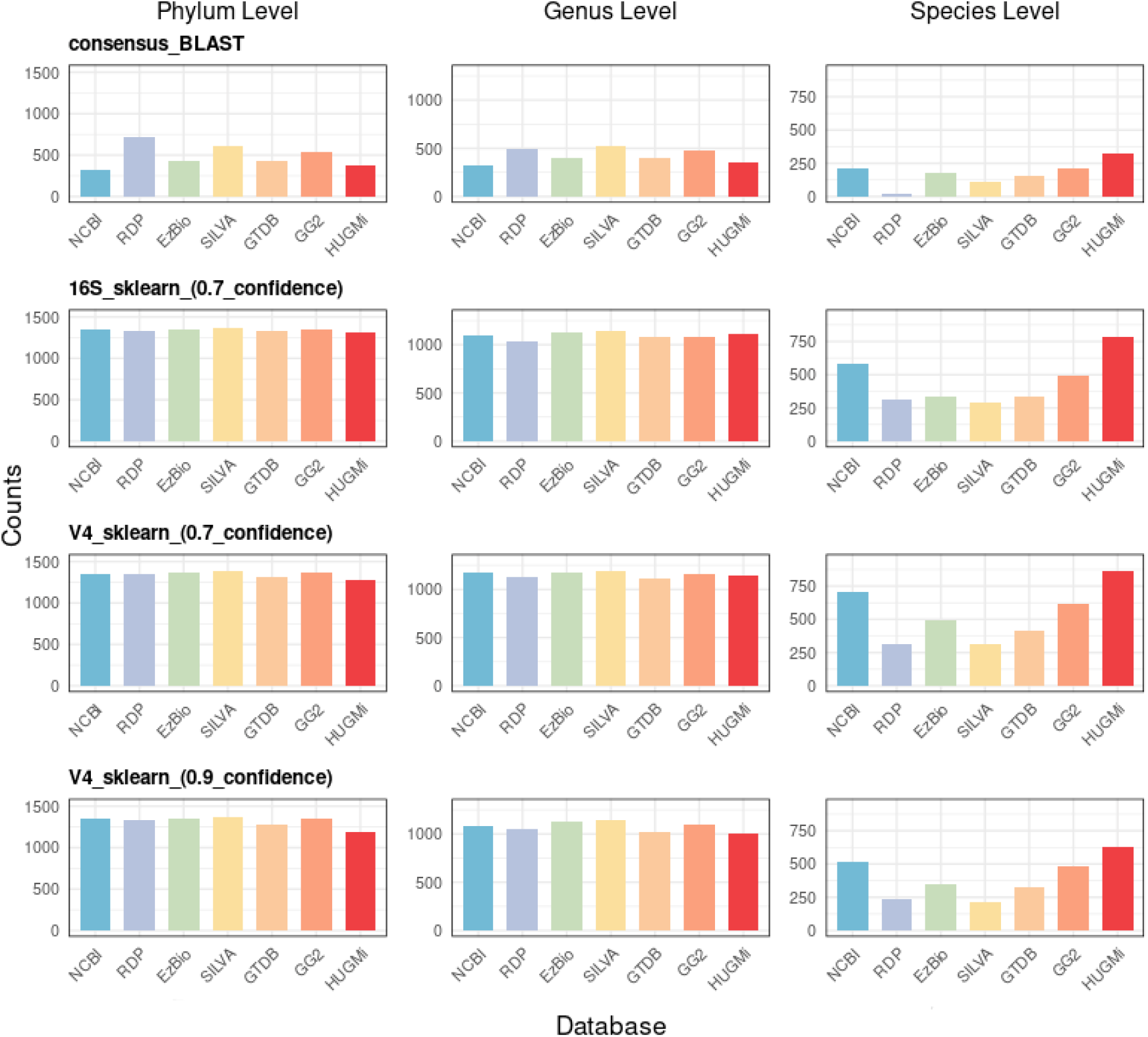
Illustration of taxonomic assignment counts at Phylum, Genus and Species levels across different databases and classifier conditions on dataset I.

#### Exploring taxonomic assignment capabilities of HUGMi database paired with QIIME2 sklearn classifier

Although BLAST methods deliver specificity with the assigned taxa, the sklearn (Bokulich et al., 2018) classifier based method is more ubiquitously used nowadays, due to its robustness, speed and posterior probability approach. Hence the performance of the HUGMi database with this learning-based was also inquired. To evaluate the full spectrum of taxonomic changes here, stringent confidence cutoffs (0.8 and 0.9) were used alongside the default (0.7), and whole 16S classifiers as well as region specific classifiers were also employed. The results for dataset I using QIIME2 sklearn classifier at varying parameters and conditions are demonstrated in Figure 3, and it confirms that HUGMi gives notably higher number of species level assignments, while being at par with the rest of the database at phylum and genus levels. In order to replicate the findings, the same classifiers with varying conditions and parameters were implemented on datasets II and III. Detailed outcomes of all the runs are presented in Supplementary Tables 6, 7 and 8.

### Taxonomic classification improvements achieved through a novel integrated bioinformatics approach

HUGMi database implemented with QIIME2 feature classifiers has already demonstrated a higher yield of species-level taxonomic assignment with both mock communities and real time clinical data. Improving upon it, the HUGMi hybrid classifier unifies the advantages of both, BLAST and sklearn-based classifiers. Figure 4 elucidates that the hybrid classifier assigns a greater number of species assignments on both, datasets I and II, as compared to its counterparts. Even with changing values of classifier confidence, or 16S regions, this hybrid classifier yields a better result (Supplementary Table 9).

**Figure 4:**
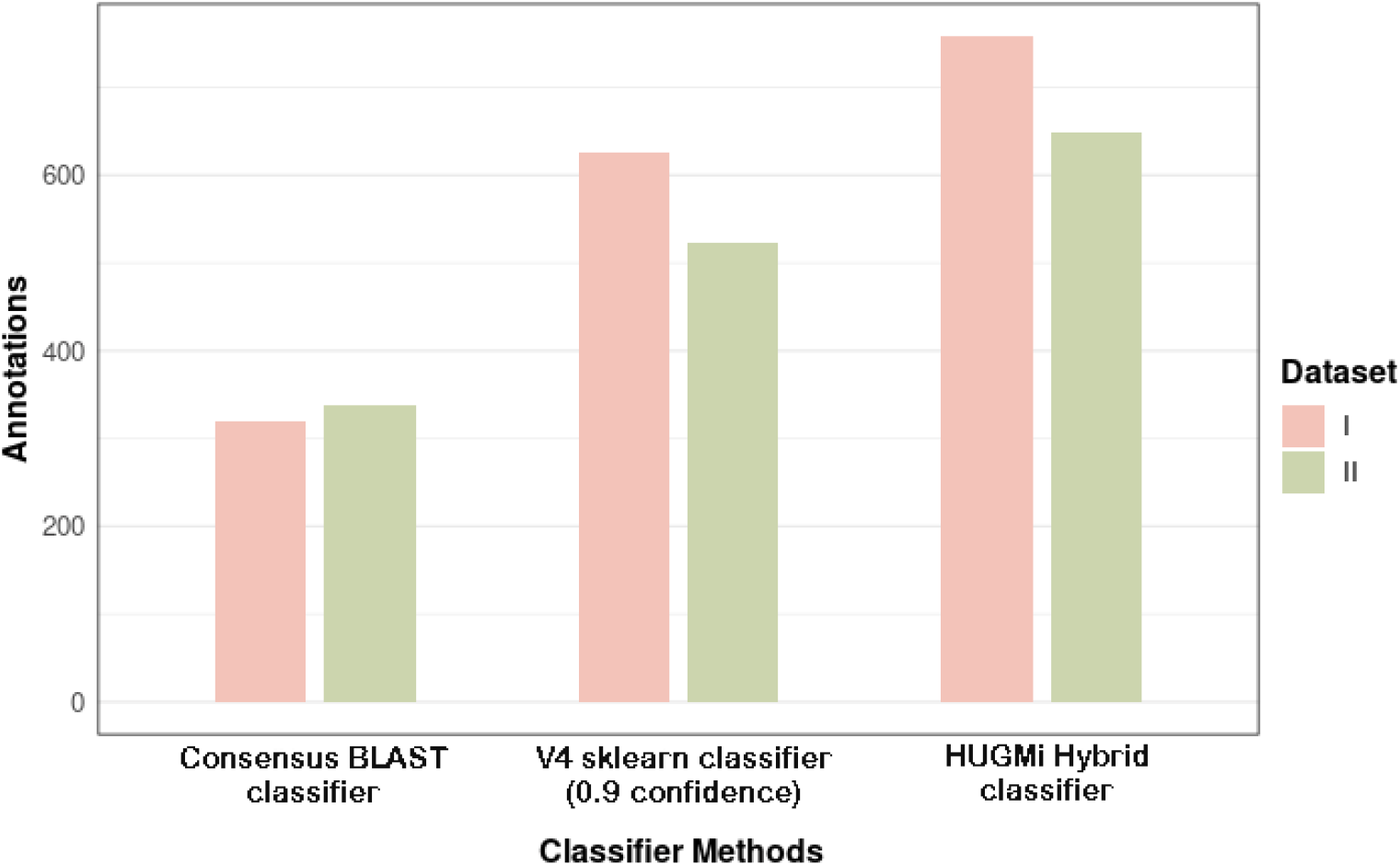
Boxplot for taxonomic annotation counts for datasets I and II using different classifiers.

### Comparative analysis of taxonomic assignments between the source studies and HUGMi framework

The taxonomic assignments by the HUGMi database along with the hybrid classifier (V4, 0.9 confidence) for datasets I and II, were compared with the original findings of the respective studies. For dataset I, as demonstrated in Supplementary Table 10, the number of taxonomies assigned in the study is far lower than the overall assignments by HUGMi database and hybrid classifier. At the phylum-level, 7 out of the 9 reported taxa matched, however HUGMi provided additional 4 phylum assignments. At genus-level, 62 genera were reported by the study whereas HUGMi predicted 156 genera, out of which 53 overlapped. Finally at the species-level, 7 bacterial names were reported, out of which 1 was mislabeled. Among the 6 valid designations, HUGMi successfully recognized 5 while simultaneously detecting an additional 252 species not documented in the initial investigation (Figure 5).

**Figure 5:**
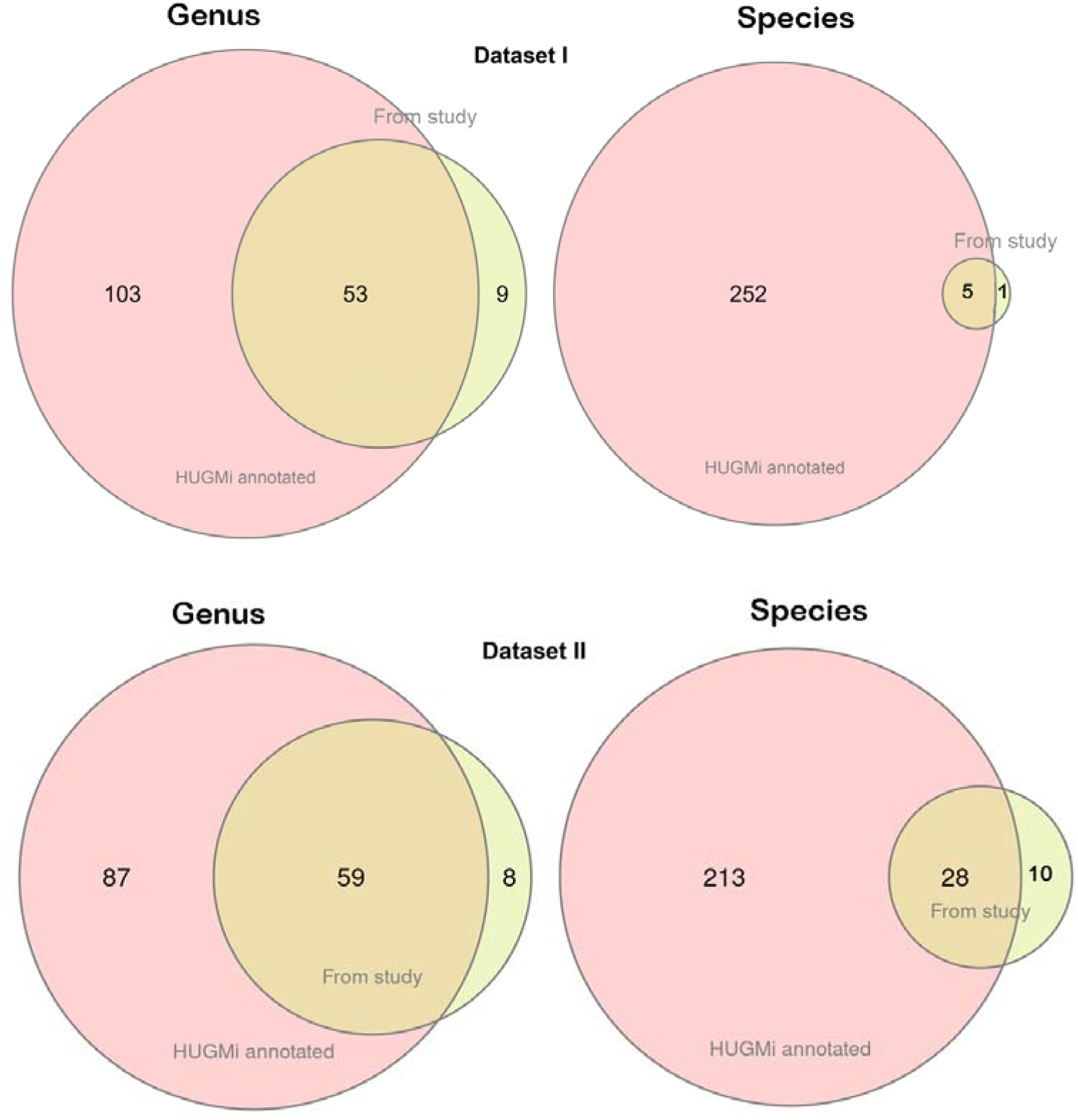
Venn diagram depicting overlap between HUGMi annotations and observations from the respective studies.

The taxonomy comparison from dataset II was done in two ways (Supplementary Table 11). The first set of comparison was the observed taxa in the study, where out of the 7 genera and 7 species identified, 6 genera and 5 species overlapped with HUGMi assignment. In the second set of comparison, the taxonomies from closed-reference OTU picking were compared with the of HUGMi findings and shown in Figure 5. Out of the 67 genera, HUGMi found 59, along with 87 more, while at species-level 28 overlapped, and HUGMi additionally observed 213 more bacterial species.

### Biological implications of HUGMi framework

Differential analysis of both validation datasets after annotation through the HUGMi framework identified an elevated number of species-level identification associated with the respective groups. Analysing through the statistically significant taxa among them revealed few biologically relevant species which remained undetected in the studies. In case of dataset I, which dealt with bladder cancer, *Streptococcus agalactiae* and *Escherichia coli* were found to be differentially significant in the cancer group, while *Gemella haemolysans* was in the healthy group. *S. agalactiae* has a well known association with recurrent urinary tract infection (UTI), which increases the risk of bladder cancer (Bayne et al., 2018; Mohanty et al., 2021). Even uropathogenic *E. coli*, which is not only associated with UTI but also produces cytotoxic necrotizing factor 1 (CNF1), a toxin known to promote bladder cancer angiogenesis (Guo et al., 2020). On the contrary, *G. haemolysans* is a noted commensal of the human genitourinary tract (Khan et al., 2004).

In the second dataset, *Sneathia sanguinegens* was found to be significant in the cervical cancer group, while *Lactobacillus jensenii* was significant for the healthy group. *S. sanguinegens* has been prominently associated with high-grade squamous intraepithelial lesions (Mitra et al., 2015), which progresses to cervical cancer. In contrast, *L. jensenii* is classically found in a healthy cervical environment and is one the dominant members of the community-state type V (CST-V) (Ma et al., 2012; Ravel et al., 2011) (Supplementary Table 12).

## DISCUSSION

Microbiome studies with the traditional microbe presence-absence layout are imminently transitioning to functional level study, and a pivotal component for that is microbial identification at a finer level (e.g., genus, or species). The HUGMi database attempts to achieve that using the cost-efficient amplicon sequencing techniques. Its key defining attributes are i) improved species-level identification, ii) niche specificity and biological relevance, and iii) nomenclature consistency. To achieve uniform taxonomic changes at all seven levels, bacterial names were updated according to the International Committee on Systematics of Prokaryotes (ICSP) nomenclature alterations from 2022 till 2024. The HUGMi database has allocated all renamed phylums, classes, as well as the segregation of many genera including *Lactobacillus* and *Prevotella*, renaming of genus for *Atopobium vaginae* to *Fannyhessea vaginae* and even certain changes in the orthography of various species. Alongside the taxonomic revision, HUGMi provides a repertoire of 1017 bacterial species, which are exclusive to the human urogenital region; unlike gut, microbiome of this particular domain remains largely unexplored and the HUGMi database might contribute to that end.

Our empirical observations reveal a more refined performance of HUGMi with respect to other databases, in its ability to assign species-level taxonomy from amplicon sequencing data. The quantitative benchmarking of HUGMi database leveraging mock communities revealed that it has consistently outperformed in precision and overall accuracy, resulting in statistically significant difference in the mean F1 scores (Supplementary Table 4). This corroborates the overall reliable performance of HUGMi database in annotation of bacterial species. The efficacy of HUGMi transcends mock communities and it has demonstrated to perform reliably well in real-world clinical samples of microbiome. Three different sets of real-time data were considered in the present study - dataset I (Popović et al. (2017) (Bučević Popović et al., 2018)), dataset II (Ilhan et al. (2019) (Ilhan et al., 2019)) and dataset III (Musa et al. (2023) (Musa et al., 2023)). The first two datasets were different kinds of human samples from the urogenital region (urine and vaginal swab, respectively) of patients suffering from bladder cancer and cervical cancer respectively. Since these two datasets were from the same 16S region (V4), a third cervicovaginal dataset that explored a different 16S region (V3-V4) was incorporated and the results are shown in Supplementary Table 8.

With the QIIME2 BLAST classifier (Camacho et al., 2009) having stringent identity and query-coverage parameters, HUGMi showed a robust performance. While deploying QIIME2 learning-based (sklearn) classifiers (Bokulich et al., 2018) across varying confidence thresholds (0.7, 0.8, and 0.9) and different 16S regions, it replicated a similar level of performance. In the case of the BLAST method, it gave comparable assignments at phylum and genus levels with that of the other databases, while it substantially improved at the species level. This enhanced resolution at the species level might be particularly helpful for researchers to derive ecological roles of microbes in viable complex communities. HUGMi exhibited a similar, consistent pattern of assignment in case of QIIME2 sklearn classifier, across multiple parameters. This highlights the versatility of HUGMi in handling a diverse set of analytical scenarios.

These results paved the way for designing a novel HUGMi hybrid classifier, which integrates the strengths of both BLAST and sklearn-based approaches. It consolidates specificity of BLAST-based methods, as well as the robustness and probabilistic framework of machine learning classifiers, resulting in superior species-level identification despite the varying confidence thresholds. The hybrid classifier coupled with the HUGMi database will likely encourage microbiome studies of various urogenital pathophysiologies and lead to a better understanding of disease progression in these body sites.

Lastly the output of the HUGMi framework on datasets I and II were compared with the outcomes of the actual studies. This helped in the evaluation of HUGMi performance on true biological data. In both cases there was major overlap in the taxa found in the studies and annotated by HUGMi. Although, in case of HUGMi, there are a large number of additional genera and species identification. The few taxa that were not identified by HUGMi could be due to the fact that studies used earlier versions of databases which might have different sets of sequences and annotations from that of HUGMi. Another explanation was that genus like *Helcococcus* and *Kocuria* in both datasets I and II, is actually not present in the current HMP list of urogenital microbes, thus not included in HUGMi. Nonetheless, the differential analysis underscores the importance of enhanced taxonomic resolution, where the newly identified bacterial species might elucidate the functional contributions in urogenital diseases.

Alongside all its benefits, HUGMi database possess the limitations of a secondary database, which includes i) dependance on primary database to ensure sequence quality and incorporation of novel sequences, ii) it requires a regular updation with the publication of higher versions of primary database, iii) identification of novel species will require a more comprehensive database and iv) it is limited to a particular human body site microbiota. However the proposed methodology of creating HUGMi, that has been described here, can be used as a generalized protocol for creating any body site specific secondary database, which might aid in higher resolution of taxonomic assignment in those niches, from amplicon sequencing.

## FUTURE DIRECTION

As urogenital research progresses, novel bacterial species might get identified. The aim is to constantly update the bacterial repertoire and the database itself by including those taxa. Also, as newer versions of the constituent databases are published, HUGMi upgradation would be necessary.

## AUTHOR DECLARATION

The authors declare no conflict of interest.

## DATA AVAILABILITY

The HUGMi database and hybrid classifier is available at https://github.com/debaleena-bhowmik/HUGMi. The accession numbers of all the validation datasets are available in Supplementary Table 3.

## ACKNOWLEDGEMENT

SP acknowledges support from SERB extramural fund CRG/2022/005998. DB acknowledges Indian Council of Medical Research (ICMR) for providing fellowship; Mr. Abhishake Lahiri for valuable discussions.

## Notes

### Competing Interest Statement

The authors have declared no competing interest.

## REFERENCES

Amir, A., McDonald, D., Navas-Molina, J. A., Kopylova, E., Morton, J. T., Zech Xu, Z., Kightley, E. P., Thompson, L. R., Hyde, E. R., Gonzalez, A., & Knight, R. (2017). Deblur Rapidly Resolves Single-Nucleotide Community Sequence Patterns. mSystems, 2(2), 10.1128/msystems.00191-16. 10.1128/msystems.00191-16

Bayne, C. E., Farah, D., Herbst, K. W., & Hsieh, M. H. (2018). Role of urinary tract infection in bladder cancer: A systematic review and meta-analysis. World Journal of Urology, 36(8), 1181–1190. 10.1007/s00345-018-2257-z

Bedford, L., Parker, S. E., Davis, E., Salzman, E., Hillier, S. L., Foxman, B., & Harlow, B. L. (2020). Characteristics of the vaginal microbiome in women with and without clinically confirmed vulvodynia. American Journal of Obstetrics and Gynecology, 223(3), 406.e1-406.e16. 10.1016/j.ajog.2020.02.039

Bhide, A., Tailor, V., & Khullar, V. (2020). Interstitial cystitis/bladder pain syndrome and recurrent urinary tract infection and the potential role of the urinary microbiome. Post Reproductive Health, 26(2), 87–90. 10.1177/2053369120936426

Bokulich, N. A., Kaehler, B. D., Rideout, J. R., Dillon, M., Bolyen, E., Knight, R., Huttley, G. A., & Gregory Caporaso, J. (2018). Optimizing taxonomic classification of marker-gene amplicon sequences with QIIME 2’s q2-feature-classifier plugin. Microbiome, 6(1), 90. 10.1186/s40168-018-0470-z

Bolyen, E., Rideout, J. R., Dillon, M. R., Bokulich, N. A., Abnet, C. C., Al-Ghalith, G. A., Alexander, H., Alm, E. J., Arumugam, M., Asnicar, F., Bai, Y., Bisanz, J. E., Bittinger, K., Brejnrod, A., Brislawn, C. J., Brown, C. T., Callahan, B. J., Caraballo-Rodríguez, A. M., Chase, J., … Caporaso, J. G. (2019). Reproducible, interactive, scalable and extensible microbiome data science using QIIME 2. Nature Biotechnology, 37(8), pArticle 8. 10.1038/s41587-019-0209-9

Bučević Popović, V., Šitum, M., Chow, C.-E. T., Chan, L. S., Roje, B., & Terzić, J. (2018). The urinary microbiome associated with bladder cancer. Scientific Reports, 8(1), 12157. 10.1038/s41598-018-29054-w

Callahan, B. J., McMurdie, P. J., Rosen, M. J., Han, A. W., Johnson, A. J. A., & Holmes, S. P. (2016). DADA2: High-resolution sample inference from Illumina amplicon data. Nature Methods, 13(7), 581–583. 10.1038/nmeth.3869

Camacho, C., Coulouris, G., Avagyan, V., Ma, N., Papadopoulos, J., Bealer, K., & Madden, T. L. (2009). BLAST+: Architecture and applications. BMC Bioinformatics, 10(1), 421. 10.1186/1471-2105-10-421

Chen, T., Yu, W.-H., Izard, J., Baranova, O. V., Lakshmanan, A., & Dewhirst, F. E. (2010). The Human Oral Microbiome Database: A web accessible resource for investigating oral microbe taxonomic and genomic information. Database: The Journal of Biological Databases and Curation, 2010, baq013. 10.1093/database/baq013

Cole, J. R., Wang, Q., Fish, J. A., Chai, B., McGarrell, D. M., Sun, Y., Brown, C. T., Porras-Alfaro, A., Kuske, C. R., & Tiedje, J. M. (2014). Ribosomal Database Project: Data and tools for high throughput rRNA analysis. Nucleic Acids Research, 42(Database issue), D633–D642. 10.1093/nar/gkt1244

DeSantis, T. Z., Hugenholtz, P., Larsen, N., Rojas, M., Brodie, E. L., Keller, K., Huber, T., Dalevi, D., Hu, P., & Andersen, G. L. (2006). Greengenes, a chimera-checked 16S rRNA gene database and workbench compatible with ARB. Applied and Environmental Microbiology, 72(7), 5069–5072. 10.1128/AEM.03006-05

Dueholm, M. S., Andersen, K. S., McIlroy, S. J., Kristensen, J. M., Yashiro, E., Karst, S. M., Albertsen, M., & Nielsen, P. H. (2020). Generation of Comprehensive Ecosystem-Specific Reference Databases with Species-Level Resolution by High-Throughput Full-Length 16S rRNA Gene Sequencing and Automated Taxonomy Assignment (AutoTax). mBio, 11(5), 10.1128/mbio.01557-20. 10.1128/mbio.01557-20

Edgar, R. C. (2010). Search and clustering orders of magnitude faster than BLAST. Bioinformatics, 26(19), 2460–2461. 10.1093/bioinformatics/btq461

Edgar, R. C. (2016). UNOISE2: Improved error-correction for Illumina 16S and ITS amplicon sequencing (p. 081257). bioRxiv. 10.1101/081257

Guo, Y., Wang, J., Zhou, K., Lv, J., Wang, L., Gao, S., Keller, E. T., Zhang, Z.-S., Wang, Q., & Yao, Z. (2020). Cytotoxic necrotizing factor 1 promotes bladder cancer angiogenesis through activating RhoC. The FASEB Journal, 34(6), 7927–7940. 10.1096/fj.201903266RR

Holm, J. B., Gajer, P., & Ravel, J. (2024). SpeciateIT and vSpeciateDB: Novel, fast and accurate per sequence 16S rRNA gene taxonomic classification of vaginal microbiota. bioRxiv: The Preprint Server for Biology, 2024.04.18.590089. 10.1101/2024.04.18.590089

Ii, M. S. R., O’Rourke, D. R., Kaehler, B. D., Ziemski, M., Dillon, M. R., Foster, J. T., & Bokulich, N. A. (2021). RESCRIPt: Reproducible sequence taxonomy reference database management. PLOS Computational Biology, 17(11), e1009581. 10.1371/journal.pcbi.1009581

Ilhan, Z. E., Łaniewski, P., Thomas, N., Roe, D. J., Chase, D. M., & Herbst-Kralovetz, M. M. (2019). Deciphering the complex interplay between microbiota, HPV, inflammation and cancer through cervicovaginal metabolic profiling. EBioMedicine, 44, 675–690. 10.1016/j.ebiom.2019.04.028

Khan, R., Urban Carl,, Rubin David,, & and Segal-maurer, S. (2004). Subacute endocarditis caused by Gemella haemolysans and a review of the literature. Scandinavian Journal of Infectious Diseases, 36(11–12), 885–888. 10.1080/00365540410024916

Love, M. I., Huber, W., & Anders, S. (2014). Moderated estimation of fold change and dispersion for RNA-seq data with DESeq2. Genome Biology, 15(12), 550. 10.1186/s13059-014-0550-8

Ma, B., Forney, L. J., & Ravel, J. (2012). The vaginal microbiome: Rethinking health and diseases. Annual Review of Microbiology, 66, 371–389. 10.1146/annurev-micro-092611-150157

Martiny, J. B. H., Jones, S. E., Lennon, J. T., & Martiny, A. C. (2015). Microbiomes in light of traits: A phylogenetic perspective. Science, 350(6261), aac9323. 10.1126/science.aac9323

McDonald, D., Jiang, Y., Balaban, M., Cantrell, K., Zhu, Q., Gonzalez, A., Morton, J. T., Nicolaou, G., Parks, D. H., Karst, S. M., Albertsen, M., Hugenholtz, P., DeSantis, T., Song, S. J., Bartko, A., Havulinna, A. S., Jousilahti, P., Cheng, S., Inouye, M., … Knight, R. (2024). Greengenes2 unifies microbial data in a single reference tree. Nature Biotechnology, 42(5), 715–718. 10.1038/s41587-023-01845-1

McIlroy, S. J., Saunders, A. M., Albertsen, M., Nierychlo, M., McIlroy, B., Hansen, A. A., Karst, S. M., Nielsen, J. L., & Nielsen, P. H. (2015). MiDAS: The field guide to the microbes of activated sludge. Database, 2015, bav062. 10.1093/database/bav062

McMurdie, P. J., & Holmes, S. (2013). phyloseq: An R Package for Reproducible Interactive Analysis and Graphics of Microbiome Census Data. PLOS ONE, 8(4), e61217. 10.1371/journal.pone.0061217

Mikaelyan, A., Köhler, T., Lampert, N., Rohland, J., Boga, H., Meuser, K., & Brune, A. (2015). Classifying the bacterial gut microbiota of termites and cockroaches: A curated phylogenetic reference database (DictDb). Systematic and Applied Microbiology, 38(7), 472–482. 10.1016/j.syapm.2015.07.004

Mitra, A., MacIntyre, D. A., Lee, Y. S., Smith, A., Marchesi, J. R., Lehne, B., Bhatia, R., Lyons, D., Paraskevaidis, E., Li, J. V., Holmes, E., Nicholson, J. K., Bennett, P. R., & Kyrgiou, M. (2015). Cervical intraepithelial neoplasia disease progression is associated with increased vaginal microbiome diversity. Scientific Reports, 5, 16865. 10.1038/srep16865

Mohanty, S., Purohit, G., Rath, S., Seth, R. K., & Mohanty, R. R. (2021). Urinary tract infection due to Group B Streptococcus: A case series from Eastern India. Clinical Case Reports, 9(10), e04885. 10.1002/ccr3.4885

Molano, L.-A. G., Vega-Abellaneda, S., & Manichanh, C. (2024). GSR-DB: A manually curated and optimized taxonomical database for 16S rRNA amplicon analysis. mSystems, 9(2), e00950–23. 10.1128/msystems.00950-23

Musa, J., Maiga, M., Green, S. J., Magaji, F. A., Maryam, A. J., Okolo, M., Nyam, C. J., Cosmas, N. T., Silas, O. A., Imade, G. E., Zheng, Y., Joyce, B. T., Diakite, B., Morhason-Bello, I., Achenbach, C. J., Sagay, A. S., Ujah, I. A. O., Murphy, R. L., Hou, L., & Mehta, S. D. (2023). Vaginal microbiome community state types and high-risk human papillomaviruses in cervical precancer and cancer in North-central Nigeria. BMC Cancer, 23(1), 683. 10.1186/s12885-023-11187-5

O’Leary, N. A., Wright, M. W., Brister, J. R., Ciufo, S., Haddad, D., McVeigh, R., Rajput, B., Robbertse, B., Smith-White, B., Ako-Adjei, D., Astashyn, A., Badretdin, A., Bao, Y., Blinkova, O., Brover, V., Chetvernin, V., Choi, J., Cox, E., Ermolaeva, O., … Pruitt, K. D. (2016). Reference sequence (RefSeq) database at NCBI: Current status, taxonomic expansion, and functional annotation. Nucleic Acids Research, 44(D1), D733–745. 10.1093/nar/gkv1189

Parks, D. H., Chuvochina, M., Rinke, C., Mussig, A. J., Chaumeil, P.-A., & Hugenholtz, P. (2022). GTDB: An ongoing census of bacterial and archaeal diversity through a phylogenetically consistent, rank normalized and complete genome-based taxonomy. Nucleic Acids Research, 50(D1), D785–D794. 10.1093/nar/gkab776

Perez-Carrasco, V., Soriano-Lerma, A., Soriano, M., Gutiérrez-Fernández, J., & Garcia-Salcedo, J. A. (2021). Urinary Microbiome: Yin and Yang of the Urinary Tract. Frontiers in Cellular and Infection Microbiology, 11. 10.3389/fcimb.2021.617002

Pernigoni, N., Guo, C., Gallagher, L., Yuan, W., Colucci, M., Troiani, M., Liu, L., Maraccani, L., Guccini, I., Migliorini, D., de Bono, J., & Alimonti, A. (2023). The potential role of the microbiota in prostate cancer pathogenesis and treatment. Nature Reviews Urology, 20(12), 706–718. 10.1038/s41585-023-00795-2

Quast, C., Pruesse, E., Yilmaz, P., Gerken, J., Schweer, T., Yarza, P., Peplies, J., & Glöckner, F. O. (2013). The SILVA ribosomal RNA gene database project: Improved data processing and web-based tools. Nucleic Acids Research, 41(Database issue), D590– D596. 10.1093/nar/gks1219

Ravel, J., Gajer, P., Abdo, Z., Schneider, G. M., Koenig, S. S. K., McCulle, S. L., Karlebach, S., Gorle, R., Russell, J., Tacket, C. O., Brotman, R. M., Davis, C. C., Ault, K., Peralta, L., & Forney, L. J. (2011). Vaginal microbiome of reproductive-age women. Proceedings of the National Academy of Sciences of the United States of America, 108(Suppl 1), 4680–4687. 10.1073/pnas.1002611107

Ritari, J., Salojärvi, J., Lahti, L., & de Vos, W. M. (2015). Improved taxonomic assignment of human intestinal 16S rRNA sequences by a dedicated reference database. BMC Genomics, 16(1), 1056. 10.1186/s12864-015-2265-y

Rohwer, R. R., Hamilton, J. J., Newton, R. J., & McMahon, K. D. (2018). TaxAss: Leveraging a Custom Freshwater Database Achieves Fine-Scale Taxonomic Resolution. mSphere, 3(5), 10.1128/msphere.00327-18. 10.1128/msphere.00327-18

Sayers, E. (2022). A General Introduction to the E-utilities. In Entrez Programming Utilities Help [Internet]. National Center for Biotechnology Information (US). https://www.ncbi.nlm.nih.gov/books/NBK25497/

Seedorf, H., Kittelmann, S., Henderson, G., & Janssen, P. H. (2014). RIM-DB: A taxonomic framework for community structure analysis of methanogenic archaea from the rumen and other intestinal environments. PeerJ, 2, e494. 10.7717/peerj.494

Steinegger, M., & Söding, J. (2017). MMseqs2 enables sensitive protein sequence searching for the analysis of massive data sets. Nature Biotechnology, 35(11), 1026–1028. 10.1038/nbt.3988

Thomas-White, K., Forster, S. C., Kumar, N., Van Kuiken, M., Putonti, C., Stares, M. D., Hilt, E. E., Price, T. K., Wolfe, A. J., & Lawley, T. D. (2018). Culturing of female bladder bacteria reveals an interconnected urogenital microbiota. Nature Communications, 9(1), 1557. 10.1038/s41467-018-03968-5

Turnbaugh, P. J., Ley, R. E., Hamady, M., Fraser-Liggett, C. M., Knight, R., & Gordon, J. I. (2007). The human microbiome project. Nature, 449(7164), 804–810. 10.1038/nature06244

Xu, H., Tamrat, N. E., Gao, J., Xu, J., Zhou, Y., Zhang, S., Chen, Z., Shao, Y., Ding, L., Shen, B., & Wei, Z. (2021). Combined Signature of the Urinary Microbiome and Metabolome in Patients With Interstitial Cystitis. Frontiers in Cellular and Infection Microbiology, 11, 711746. 10.3389/fcimb.2021.711746

Yoon, S.-H., Ha, S.-M., Kwon, S., Lim, J., Kim, Y., Seo, H., & Chun, J. (2017). Introducing EzBioCloud: A taxonomically united database of 16S rRNA gene sequences and whole-genome assemblies. International Journal of Systematic and Evolutionary Microbiology, 67(5), 1613–1617. 10.1099/ijsem.0.001755

