## Supplementary Table 2 for "HUGMi: Human Uro-Genital Microbiome database and hybrid classifier for improved species level annotation of 16S rRNA amplicon sequences"

**Supplementary Table 2: MeSH searches in NCBI for relevant papers**

| MeSH Terms | No. of papers |
|---|---|
| ((("Urinary Tract"[MeSH] OR "Urinary Bladder"[MeSH] OR "Urine"[MeSH]) AND ("Microbiota"[MeSH] OR "Microbiome"[TW] OR "Bacteria"[MeSH])) AND ("Health"[MeSH] OR "Urologic Diseases"[MeSH] OR "Interstitial Cystitis"[TW] OR "Bladder Cancer"[TW] OR "Kidney Failure"[TW] OR "Urolithiasis"[MeSH] OR "Urinary Tract Infections"[MeSH] OR "Prostitis"[TW] OR "Hematuria"[MeSH] OR "Acquired Immunodeficiency Syndrome"[MeSH] OR "Benign Prostatic Hypertrophy"[TW] OR "Benign Prostatic Hyperplasia"[TW] OR "Prostate Cancer"[TW] OR "Vesicocutaneous Fistula"[TW] OR "Urinary Incontinence"[MeSH] OR "Pelvic Floor Disorders"[MeSH] OR "Pylonephritis"[TW] OR "Cystocele"[MeSH] OR "Urethritis"[MeSH] OR "Sexually Transmitted Diseases"[MeSH])) AND ("infant"[MeSH] OR "child"[MeSH] OR "adolescent"[MeSH] OR "adult"[MeSH]) AND ("last 5 years"[PDat]) | 203 |
| ((("Vagina"[MeSH] OR "Vaginal Smear"[TW]) AND ("Microbiota"[MeSH] OR "Microbiome"[TW] OR "Bacteria"[MeSH])) AND ("Health"[MeSH] OR "Vaginal Diseases"[MeSH] OR "Bacterial Vaginosis"[TW] OR "Yeast Infection"[TW] OR "Gonorrhoea"[TW] OR "Trichomoniasis"[TW] OR "Human Papillomavirus"[TW] OR "Pregnant"[TW] OR "Acquired Immunodeficiency Syndrome"[MeSH] OR "Sexually Transmitted Diseases"[MeSH])) AND ("infant"[MeSH] OR "child"[MeSH] OR "adolescent"[MeSH] OR "adult"[MeSH]) AND ("last 5 years"[PDat]) | 368 |
| ((("Penis"[MeSH] OR "Penile Skin"[TW] OR "Penile Discharge"[TW]) AND ("Microbiota"[MeSH] OR "Microbiome"[TW] OR "Bacteria"[MeSH])) AND ("Health"[MeSH] OR "Penile Diseases"[MeSH] OR "Balanitis"[MeSH] OR "Bowen's Disease"[MeSH] OR "Erectile dysfunction"[MeSH] OR "Genital Wart"[TW] OR "Lichen Planus"[MeSH] OR "Lichen sclerosus"[TW] OR "Penile Cancer"[TW] OR "Priapism"[MeSH] OR "Acquired Immunodeficiency Syndrome"[MeSH] OR "Sexually Transmitted Diseases"[MeSH])) AND ("infant"[MeSH] OR "child"[MeSH] OR "adolescent"[MeSH] OR "adult"[MeSH]) AND ("last 5 years"[PDat]) | 32 |
