## Supplementary Table 3 for "HUGMi: Human Uro-Genital Microbiome database and hybrid classifier for improved species level annotation of 16S rRNA amplicon sequences"

**Supplementary Table 3: Details of real time datasets**

| Dataset | Accession no. | Disease | Sample type | 16S region | Sequencing platform |
|---|---|---|---|---|---|
| I | PRJEB22327 | Badder cancer | urine | V4 | Illumina |
| II | PRJNA518153 | Cervical cancer and HPV infection | vaginal swab | V4 | Illumina |
| III | PRJNA914231 | Cervical cancer and precancer states | cervicovaginal lavage | V3V4 | 454 pyrosequnecing |
