## Supplementary Table 4 for "HUGMi: Human Uro-Genital Microbiome database and hybrid classifier for improved species level annotation of 16S rRNA amplicon sequences"

**Supplementary Table 4: t-test between HUGMi and each of the database scores**

**Genus-level**

| Database name | p value |
|---|---|
| EzBio | 2.45E-07 |
| GG2 | 9.26E-04 |
| GTDB | 1.65E-07 |
| NCBI | 1.90E-08 |
| SILVA | 1.82E-03 |
| RDP | 3.61E-05 |

**Species-level**

| Database name | p value |
|---|---|
| EzBio | 5.51E-09 |
| GG2 | 3.56E-08 |
| GTDB | 7.37E-09 |
| NCBI | 8.55E-09 |
| SILVA | 6.52E-08 |
| RDP | 9.90E-10 |
