## Supplementary Table 5 for "HUGMi: Human Uro-Genital Microbiome database and hybrid classifier for improved species level annotation of 16S rRNA amplicon sequences"

### Supplementary Table 5: Taxonomic assignment counts of dataset I

**BLAST parameters: perc-identity = 100%; query-cov = 95%; max-accepts = 10)**

| Database | domain | phylum | class | order | family | genus | species |
|---|---|---|---|---|---|---|---|
| EzBio | 435 | 435 | 435 | 434 | 431 | 395 | 177 |
| GG2 | 539 | 537 | 537 | 530 | 525 | 472 | 217 |
| GTDB | 425 | 422 | 422 | 421 | 417 | 401 | 159 |
| NCBI | 322 | 322 | 322 | 322 | 320 | 315 | 213 |
| RDP | 726 | 721 | 514 | 511 | 511 | 499 | 21 |
| SILVA | 602 | 602 | 601 | 596 | 582 | 521 | 108 |
| HUGMi | 386 | 376 | 371 | 365 | 363 | 358 | 320 |
