## Supplementary Table 6 for "HUGMi: Human Uro-Genital Microbiome database and hybrid classifier for improved species level annotation of 16S rRNA amplicon sequences"

**Supplementary Table 6: Taxonomic assignment counts of dataset I using sklearn**

**confidence 0.7 (default)**

**16S**

| Database | domain | phylum | class | order | family | genus | species |
|---|---|---|---|---|---|---|---|
| EzBio | 1504 | 1357 | 1349 | 1342 | 1324 | 1133 | 341 |
| GG2 | 1396 | 1343 | 1339 | 1308 | 1262 | 1086 | 498 |
| GTDB | 1566 | 1322 | 1317 | 1289 | 1242 | 1079 | 334 |
| NCBI | 1527 | 1348 | 1303 | 1282 | 1225 | 1089 | 585 |
| RDP | 1556 | 1324 | 1104 | 1100 | 1093 | 1041 | 318 |
| SILVA | 1379 | 1371 | 1369 | 1345 | 1321 | 1147 | 287 |
| HUGMi | 1566 | 1315 | 1277 | 1225 | 1189 | 1118 | 780 |

**V4**

| Database | domain | phylum | class | order | family | genus | species |
|---|---|---|---|---|---|---|---|
| EzBio | 1560 | 1360 | 1355 | 1350 | 1339 | 1179 | 488 |
| GG2 | 1549 | 1365 | 1365 | 1343 | 1306 | 1158 | 619 |
| GTDB | 1566 | 1318 | 1313 | 1278 | 1246 | 1113 | 417 |
| NCBI | 1539 | 1358 | 1335 | 1320 | 1273 | 1175 | 707 |
| RDP | 1564 | 1343 | 1175 | 1173 | 1166 | 1128 | 318 |
| SILVA | 1381 | 1377 | 1376 | 1367 | 1342 | 1183 | 316 |
| HUGMi | 1566 | 1272 | 1239 | 1220 | 1207 | 1146 | 858 |

**confidence 0.8**

**16S**

| Database | domain | phylum | class | order | family | genus | species |
|---|---|---|---|---|---|---|---|
| EzBio | 1504 | 1356 | 1346 | 1340 | 1317 | 1104 | 272 |
| GG2 | 1396 | 1333 | 1332 | 1297 | 1244 | 1049 | 428 |
| GTDB | 1566 | 1304 | 1301 | 1269 | 1221 | 1033 | 289 |
| NCBI | 1527 | 1337 | 1281 | 1261 | 1196 | 1050 | 507 |
| RDP | 1529 | 1325 | 1066 | 1059 | 1051 | 991 | 276 |
| SILVA | 1376 | 1369 | 1365 | 1339 | 1302 | 1119 | 254 |
| HUGMi | 1566 | 1155 | 1109 | 1076 | 1046 | 949 | 575 |

**V4**

| Database | domain | phylum | class | order | family | genus | species |
|---|---|---|---|---|---|---|---|
| EzBio | 1560 | 1360 | 1353 | 1347 | 1323 | 1159 | 434 |
| GG2 | 1549 | 1364 | 1364 | 1336 | 1305 | 1180 | 564 |
| GTDB | 1566 | 1312 | 1307 | 1272 | 1239 | 1080 | 386 |
| NCBI | 1539 | 1352 | 1327 | 1313 | 1260 | 1135 | 647 |
| RDP | 1564 | 1343 | 1155 | 1148 | 1142 | 1104 | 274 |
| SILVA | 1381 | 1376 | 1375 | 1356 | 1324 | 1154 | 292 |
| HUGMi | 1566 | 1234 | 1204 | 1184 | 1166 | 1087 | 743 |

**confidence 0.9**

**16S**

| Database | domain | phylum | class | order | family | genus | species |
|---|---|---|---|---|---|---|---|
| EzBio | 1483 | 1353 | 1337 | 1330 | 1301 | 1159 | 213 |
| GG2 | 1391 | 1319 | 1319 | 1279 | 1223 | 996 | 343 |
| GTDB | 1566 | 1275 | 1269 | 1211 | 1158 | 956 | 235 |
| NCBI | 1522 | 1324 | 1262 | 1238 | 1137 | 992 | 406 |
| RDP | 1436 | 1306 | 1012 | 1006 | 998 | 926 | 242 |
| SILVA | 1376 | 1359 | 1352 | 1319 | 1287 | 1078 | 197 |
| HUGMi | 1566 | 1064 | 1024 | 997 | 968 | 860 | 472 |

**V4**

| Database | domain | phylum | class | order | family | genus | species |
|---|---|---|---|---|---|---|---|
| EzBio | 1560 | 1358 | 1349 | 1343 | 1310 | 1123 | 348 |
| GG2 | 1548 | 1359 | 1359 | 1330 | 1287 | 1093 | 479 |
| GTDB | 1566 | 1287 | 1281 | 1245 | 1205 | 1027 | 326 |
| NCBI | 1539 | 1346 | 1306 | 1289 | 1221 | 1086 | 521 |
| RDP | 1564 | 1337 | 1121 | 1113 | 1108 | 1051 | 232 |
| SILVA | 1381 | 1375 | 1374 | 1359 | 1332 | 1138 | 218 |
| HUGMi | 1566 | 1182 | 1149 | 1126 | 1113 | 1010 | 625 |
