## Supplementary Table 7 for "HUGMi: Human Uro-Genital Microbiome database and hybrid classifier for improved species level annotation of 16S rRNA amplicon sequences"

**Supplementary Table 7: Taxonomic assignment counts of dataset II using sklearn**

**confidence 0.7 (default)**

**16S**

| Database | domain | phylum | class | order | family | genus | species |
|---|---|---|---|---|---|---|---|
| EzBio | 1148 | 1055 | 1049 | 1041 | 1021 | 870 | 307 |
| GG2 | 1060 | 1039 | 1038 | 1003 | 968 | 853 | 449 |
| GTDB | 1168 | 1027 | 1024 | 986 | 952 | 828 | 313 |
| NCBI | 1162 | 1045 | 1019 | 998 | 932 | 821 | 475 |
| RDP | 1167 | 1041 | 860 | 860 | 850 | 810 | 286 |
| SILVA | 1056 | 1054 | 1050 | 1006 | 985 | 832 | 265 |
| HUGMi | 1168 | 944 | 904 | 869 | 858 | 794 | 544 |

**V4**

| Database | domain | phylum | class | order | family | genus | species |
|---|---|---|---|---|---|---|---|
| EzBio | 1168 | 1055 | 1051 | 1043 | 1032 | 894 | 370 |
| GG2 | 1060 | 1039 | 1038 | 1003 | 968 | 853 | 449 |
| GTDB | 1168 | 1019 | 1017 | 979 | 953 | 864 | 381 |
| NCBI | 1158 | 1051 | 1033 | 1019 | 972 | 886 | 543 |
| RDP | 1168 | 1052 | 924 | 919 | 917 | 894 | 296 |
| SILVA | 1056 | 1056 | 1056 | 1030 | 1017 | 866 | 300 |
| HUGMi | 1168 | 979 | 956 | 938 | 929 | 882 | 695 |

**confidence 0.8**

**16S**

| Database | domain | phylum | class | order | family | genus | species |
|---|---|---|---|---|---|---|---|
| EzBio | 1148 | 1054 | 1044 | 1033 | 1005 | 849 | 262 |
| GG2 | 1060 | 1035 | 1035 | 996 | 959 | 830 | 410 |
| GTDB | 1168 | 1018 | 1017 | 975 | 940 | 805 | 277 |
| NCBI | 1162 | 1040 | 1000 | 978 | 899 | 788 | 419 |
| RDP | 1161 | 1030 | 839 | 837 | 832 | 790 | 251 |
| SILVA | 1056 | 1051 | 1047 | 1002 | 977 | 820 | 230 |
| HUGMi | 1168 | 880 | 847 | 825 | 811 | 728 | 451 |

**V4**

| Database | domain | phylum | class | order | family | genus | species |
|---|---|---|---|---|---|---|---|
| EzBio | 1168 | 1053 | 1049 | 1040 | 1027 | 887 | 345 |
| GG2 | 1167 | 1052 | 1052 | 1019 | 987 | 886 | 485 |
| GTDB | 1168 | 1017 | 1015 | 976 | 949 | 846 | 354 |
| NCBI | 1158 | 1050 | 1031 | 1018 | 959 | 851 | 503 |
| RDP | 1168 | 1047 | 904 | 900 | 895 | 872 | 266 |
| SILVA | 1056 | 1056 | 1054 | 1038 | 1020 | 864 | 278 |
| HUGMi | 1168 | 945 | 923 | 909 | 903 | 838 | 616 |

**confidence 0.9**

**16S**

| Database | domain | phylum | class | order | family | genus | species |
|---|---|---|---|---|---|---|---|
| EzBio | 1146 | 1054 | 1044 | 1028 | 997 | 827 | 205 |
| GG2 | 1059 | 1026 | 1026 | 990 | 946 | 792 | 344 |
| GTDB | 1168 | 995 | 993 | 937 | 895 | 751 | 225 |
| NCBI | 1160 | 1033 | 980 | 957 | 851 | 742 | 355 |
| RDP | 1126 | 1016 | 795 | 795 | 786 | 735 | 208 |
| SILVA | 1055 | 1048 | 1043 | 987 | 958 | 766 | 181 |
| HUGMi | 1168 | 833 | 811 | 790 | 778 | 674 | 381 |

**V4**

| Database | domain | phylum | class | order | family | genus | species |
|---|---|---|---|---|---|---|---|
| EzBio | 1168 | 1054 | 1045 | 1037 | 1014 | 873 | 303 |
| GG2 | 1167 | 1050 | 1050 | 1020 | 996 | 879 | 421 |
| GTDB | 1168 | 999 | 996 | 957 | 932 | 822 | 314 |
| NCBI | 1158 | 1046 | 1020 | 1003 | 920 | 808 | 427 |
| RDP | 1168 | 1042 | 882 | 882 | 879 | 828 | 228 |
| SILVA | 1056 | 1055 | 1051 | 1024 | 1000 | 830 | 249 |
| HUGMi | 1168 | 916 | 895 | 873 | 858 | 768 | 523 |
