## Supplementary Table 8 for "HUGMi: Human Uro-Genital Microbiome database and hybrid classifier for improved species level annotation of 16S rRNA amplicon sequences"

**Supplementary Table 8: Taxonomic assignment counts of dataset III using sklearn**

**confidence 0.7 (default)**

**V3V4**

| Database | domain | phylum | class | order | family | genus | species |
|---|---|---|---|---|---|---|---|
| EzBio | 21910 | 7745 | 7721 | 7662 | 7600 | 6987 | 3118 |
| GG2 | 23878 | 7848 | 7829 | 7678 | 7505 | 6873 | 4185 |
| GTDB | 25225 | 7475 | 7425 | 7298 | 7163 | 6897 | 2954 |
| NCBI | 25147 | 7728 | 7595 | 7537 | 7335 | 6881 | 4351 |
| RDP | 25213 | 7468 | 6639 | 6623 | 6585 | 6486 | 3260 |
| SILVA | 17806 | 7727 | 7714 | 7654 | 7579 | 6794 | 3159 |
| HUGMi | 25225 | 7313 | 6925 | 6771 | 6736 | 6403 | 5085 |

**confidence 0.8**

**V3V4**

| Database | domain | phylum | class | order | family | genus | species |
|---|---|---|---|---|---|---|---|
| EzBio | 23435 | 7717 | 7619 | 7584 | 7348 | 6682 | 2235 |
| GG2 | 22622 | 7541 | 7534 | 7395 | 7171 | 6001 | 2684 |
| GTDB | 25225 | 7217 | 7162 | 6892 | 6715 | 6103 | 2102 |
| NCBI | 23480 | 7648 | 7476 | 7379 | 7004 | 6148 | 3042 |
| RDP | 25093 | 6495 | 6275 | 6195 | 6020 | 5552 | 3245 |
| SILVA | 8418 | 7683 | 7643 | 7464 | 7186 | 6273 | 2217 |
| HUGMi | 25225 | 6793 | 6273 | 6042 | 5965 | 5320 | 4136 |

**confidence 0.9**

**V3V4**

| Database | domain | phylum | class | order | family | genus | species |
|---|---|---|---|---|---|---|---|
| EzBio | 23243 | 7691 | 7636 | 7572 | 7379 | 6463 | 1821 |
| GG2 | 22227 | 7472 | 7469 | 7301 | 7033 | 5699 | 2233 |
| GTDB | 25225 | 7109 | 7061 | 6759 | 6578 | 5889 | 1888 |
| NCBI | 23438 | 7599 | 7384 | 7259 | 6752 | 5841 | 2584 |
| RDP | 24778 | 6352 | 6149 | 6028 | 5797 | 5215 | 2852 |
| SILVA | 7897 | 7632 | 7584 | 7398 | 7056 | 5994 | 1726 |
| HUGMi | 25225 | 6305 | 5867 | 5564 | 5469 | 4642 | 3104 |
