## Supplementary Table 9 for "HUGMi: Human Uro-Genital Microbiome database and hybrid classifier for improved species level annotation of 16S rRNA amplicon sequences"

**Supplementary Table 9: Evaluation of HUGMi Hybrid q2 Classifier**

**blast parameters: perc-identity = 100%; query-cov = 95%; max-accepts = 10)**

| | | | | | | # taxonomic assignments | | | | | | | |
|---|---|---|---|---|---|---|---|---|---|---|---|---|---|
| **study** | **sample souce** | **disease** | **classifier region** | **classifier confidence** | **total ASVs** | **domain** | **phylum** | **class** | **order** | **family** | **genus** | **species** | **run time in sec. (40 cores)** |
| Dataset I | urine sample | blader cancer | 16S | 0.7 | 1566 | 1566 | 1330 | 1294 | 1246 | 1217 | 1161 | 889 | 57.577 |
| | | | 16S | 0.9 | | 1566 | 1094 | 1060 | 1040 | 1017 | 932 | 635 | 56.976 |
| | | | V4 | 0.7 | | 1566 | 1276 | 1244 | 1227 | 1215 | 1171 | 926 | 58.292 |
| | | | V4 | 0.9 | | 1566 | 1194 | 1165 | 1145 | 1136 | 1063 | 758 | 57.585 |
| Dataset II | vaginal swab | cervical cancer and pre-cancer states | 16S | 0.7 | 1168 | 1168 | 962 | 926 | 896 | 890 | 840 | 639 | 48.768 |
| | | | 16S | 0.9 | | 1168 | 864 | 846 | 832 | 825 | 752 | 538 | 48.748 |
| | | | V4 | 0.7 | | 1168 | 988 | 966 | 952 | 944 | 914 | 761 | 49.241 |
| | | | V4 | 0.9 | | 1168 | 936 | 918 | 899 | 891 | 835 | 649 | 49.025 |
