## Supplementary Table 10 for "HUGMi: Human Uro-Genital Microbiome database and hybrid classifier for improved species level annotation of 16S rRNA amplicon sequences"

**Supplementary Table 10: Comparison of taxonomic assignment with the actual study (dataset**

| **Phylum** | | |
|---|---|---|
| **From dataset I** | **updated names** | **HUGMi assignment** |
| Acidobacteria | Acidobacteriota | -- |
| Actinobacteria | Actinomycetota | Actinomycetota |
| Bacteroidetes | Bacteroidota | Bacteroidota |
| Cyanobacteria | Cyanobacteriota | -- |
| Firmicutes | Bacillota | Bacillota |
| Fusobacteria | Fusobacteriota | Fusobacteriota |
| Proteobacteria | Pseudomonadota | Pseudomonadota |
| Spirochaetes | Spirochaetota | Spirochaetota |
| Synergistetes | Synergistota | Synergistota |
| | | Campylobacterota |
| | | Chlamydiota |
| | | Mycoplasmatota |
| | | Thermodesulfobacteriota |

| **Genus** | | |
|---|---|---|
| **From dataset I** | **updated names** | **HUGMi assignment** |
| Acinetobacter | same | Acinetobacter |
| Actinobaculum | same | Actinobaculum |
| Actinomyces | same | Actinomyces |
| Aerococcus | same | Aerococcus |
| Aggregatibacter | same | -- |
| Alloscardovia | same | Alloscardovia |
| Anaerococcus | same | Anaerococcus |
| Arcanobacterium | same | Arcanobacterium |
| Atopobium | same | Atopobium |
| Bacteroides | same | Bacteroides |
| Bifidobacterium | same | Bifidobacterium |
| Bilophila | same | Bilophila |
| Blautia | same | Blautia |
| Brevibacterium | same | Brevibacterium |
| Campylobacter | same | Campylobacter |
| Chryseobacterium | same | Chryseobacterium |
| Clostridium | same | Clostridium |
| Collinsella | same | Collinsella |
| Comamonas | same | Comamonas |
| Coprococcus | same | -- |
| Corynebacterium | same | Corynebacterium |
| Dermabacter | same | Dermabacter |

| | | |
|---|---|---|
| Dialister | same | Dialister |
| Dorea | same | Dorea |
| Enhydrobacter | same | -- |
| Enterococcus | same | Enterococcus |
| Facklamia | same | Facklamia |
| Faecalibacterium | same | Faecalibacterium |
| Finegoldia | same | Finegoldia |
| Fusobacterium | same | Fusobacterium |
| Gallicola | same | Gallicola |
| Granulicatella | same | Granulicatella |
| Haemophilus | same | Haemophilus |
| Helcococcus | same | -- |
| Jeotgalicoccus | same | -- |
| Kocuria | same | -- |
| Lachnospira | same | Lachnospira |
| Lactobacillus | same | Lactobacillus |
| Micrococcus | same | Micrococcus |
| Mobiluncus | same | Mobiluncus |
| Mogibacterium | same | Mogibacterium |
| Moryella | same | Moryella |
| Oscillospira | same | -- |
| Parabacteroides | same | Parabacteroides |
| Paracoccus | same | Paracoccus |
| Peptococcus | same | Peptococcus |
| Peptoniphilus | same | Peptoniphilus |
| Porphyromonas | same | Porphyromonas |
| Prevotella | same | Prevotella |
| Pseudoclavibacter | same | Pseudoclavibacter |
| Pseudomonas | same | Pseudomonas |
| Pyramidobacter | same | -- |
| Rothia | same | Rothia |
| Ruminococcus | same | Ruminococcus |
| Sphingomonas | same | Sphingomonas |
| Staphylococcus | same | Staphylococcus |
| Streptobacillus | same | -- |
| Streptococcus | same | Streptococcus |
| Sutterella | same | Sutterella |
| Treponema | same | Treponema |
| Varibaculum | same | Varibaculum |
| Veillonella | same | Veillonella |
| | | Achromobacter |
| | | Acidaminococcus |
| | | Acidovorax |
| | | Actinotignum |

Aedoeadaptatus
Aeromonas
Afipia
Agathobacter
Alistipes
Alloiococcus
Alloprevotella
Altererythrobacter
Amnibacterium
Anaeroglobus
Anaerostipes
Anaerotruncus
Brachybacterium
Brevundimonas
Catonella
Clostridioides
Cutibacterium
Desulfovibrio
Dietzia
Enterobacter
Escherichia
Eubacterium
Ezakiella
Fastidiosipila
Fenollaria
Flavobacterium
Flavonifractor
Gardnerella
Gemella
Gemelliphila
Hallella
Heyndrickxia
Howardella
Hoylesella
Jonquetella
Khoudiadiopia
Kingella
Klebsiella
Lachnoanaerobaculum
Lachnoclostridium
Lancefieldella
Latilactobacillus
Lawsonella
Leclercia

Leminorella
Leptotrichia
Levilactobacillus
Ligilactobacillus
Massilia
Mediterraneibacter
Mesorhizobium
Methylobacterium
Microvirga
Moraxella
Murdochiella
Mycobacterium
Mycobacteroides
Mycolicibacterium
Mycoplasmopsis
Negativicoccus
Neisseria
Nevskia
Nosocomiicoccus
Odoribacter
Olegusella
Oligella
Paenibacillus
Pantoea
Parachlamydia
Parvimonas
Pedobacter
Pelomonas
Peptostreptococcus
Phocaeicola
Photobacterium
Propionimicrobium
Proteus
Pseudoglutamicibacter
Pseudoramibacter
Pseudoroseomonas
Rhizobium
Rhizorhabdus
Rhodanobacter
Salmonella
Schaalia
Sediminibacterium
Segatella
Shewanella

| | | |
|---|---|---|
| | | Sneathia |
| | | Solobacterium |
| | | Sphingobium |
| | | Stenotrophomonas |
| | | Ureaplasma |
| | | Variovorax |
| | | Vibrio |
| | | Weeksella |
| | | Winkia |
| | | Xanthomonas |
| | | Yersinia |

| **Species** | | |
|---|---|---|
| **From dataset I** | **updated names** | **HUGMi assignment** |
| Campylobacter hor | same | Campylobacter hominis |
| Fusobacterium nuc | same | Fusobacterium nucleatum |
| Actinobaculum mas | same | Actinobaculum massiliense |
| Jonquetella anthrop | same | Jonquetella anthropi |
| Veillonella dispar | same | Veillonella dispar |
| Streptococcus crisp | not found | -- |
| Corynebacterium a | same | -- |
| | | Achromobacter xylosoxidans |
| | | Acidaminococcus intestini |
| | | Acidaminococcus massiliensis |
| | | Acidovorax delafieldii |
| | | Acidovorax radicis |
| | | Acinetobacter baumannii |
| | | Actinomyces oris |
| | | Actinomyces provencensis |
| | | Actinotignum schaalii |
| | | Actinotignum urinale |
| | | Aedoeadaptatus coxii |
| | | Aerococcus urinae |
| | | Aerococcus viridans |
| | | Aeromonas caviae |
| | | Afipia massiliensis |
| | | Agathobacter rectalis |
| | | Alistipes finegoldii |
| | | Alistipes onderdonkii |
| | | Alistipes shahii |
| | | Alloiococcus otitis |
| | | Alloprevotella tannerae |
| | | Alloscardovia omnicolens |

Altererythrobacter epoxidivorans
Amnibacterium kyonggiense
Anaerococcus hydrogenalis
Anaerococcus lactolyticus
Anaerococcus obesiensis
Anaerococcus prevotii
Anaerococcus provencensis
Anaerococcus tetradius
Anaerococcus vaginimassiliensis
Anaeroglobus geminatus
Anaerostipes hadrus
Anaerotruncus colihominis
Arcanobacterium ihumii
Atopobium minutum
Bacteroides fragilis
Bacteroides ovatus
Bacteroides stercoris
Bacteroides uniformis
Bifidobacterium adolescentis
Bifidobacterium breve
Bifidobacterium dentium
Bifidobacterium longum
Bilophila wadsworthia
Blautia obeum
Brevibacterium equis
Brevibacterium ravenspurgense
Brevundimonas diminuta
Brevundimonas subvibrioides
Brevundimonas vesicularis
Campylobacter ureolyticus
Catonella morbi
Chryseobacterium aquaticum
Chryseobacterium indologenes
Clostridioides difficile
Clostridium butyricum
Clostridium perfringens
Collinsella aerofaciens
Comamonas testosteroni
Corynebacterium amycolatum
Corynebacterium aurimucosum
Corynebacterium diphtheriae
Corynebacterium frankenforstense
Corynebacterium genitalium
Corynebacterium glucuronolyticum

Corynebacterium jeddahense
Corynebacterium jeikeium
Corynebacterium lipophilum
Corynebacterium matruchotii
Corynebacterium minutissimum
Corynebacterium otitidis
Corynebacterium provencense
Corynebacterium pseudodiphtheriticum
Corynebacterium pseudokroppenstedtii
Corynebacterium riegelii
Cutibacterium acnes
Dermabacter hominis
Desulfovibrio desulfuricans
Desulfovibrio fairfieldensis
Dialister invisus
Dialister micraerophilus
Dialister propionicifaciens
Dietzia maris
Dorea longicatena
Enterobacter cloacae
Enterococcus cecorum
Enterococcus mundtii
Enterococcus raffinosus
Escherichia coli
Escherichia fergusonii
Eubacterium saphenum
Eubacterium yurii
Ezakiella coagulans
Ezakiella massiliensis
Facklamia languida
Faecalibacterium prausnitzii
Fastidiosipila sanguinis
Fenollaria massiliensis
Finegoldia magna
Flavobacterium antarcticum
Flavonifractor plautii
Fusobacterium gonidiaformans
Gallicola barnesae
Gardnerella vaginalis
Gemella haemolysans
Gemelliphila asaccharolytica
Granulicatella adiacens
Granulicatella elegans
Haemophilus parainfluenzae

Hallella bergensis
Hallella colorans
Heyndrickxia coagulans
Howardella ureilytica
Hoylesella nanceiensis
Hoylesella pleuritidis
Hoylesella timonensis
Khoudiadiopia massiliensis
Kingella kingae
Klebsiella aerogenes
Klebsiella oxytoca
Lachnoanaerobaculum saburreum
Lachnoclostridium urinimassiliense
Lachnospira eligens
Lactobacillus amylovorus
Lactobacillus crispatus
Lactobacillus delbrueckii
Lactobacillus fornicalis
Lactobacillus helveticus
Lactobacillus iners
Lancefieldella parvula
Latilactobacillus sakei
Lawsonella clevelandensis
Leclercia adecarboxylata
Leminorella grimontii
Leptotrichia wadei
Levilactobacillus brevis
Ligilactobacillus salivarius
Massilia timonae
Mediterraneibacter faecis
Mesorhizobium amorphae
Methylobacterium radiotolerans
Micrococcus luteus
Microvirga flocculans
Mobiluncus curtisii
Mobiluncus mulieris
Mogibacterium timidum
Moraxella nonliquefaciens
Moraxella osloensis
Moryella indoligenes
Murdochiella asaccharolytica
Murdochiella vaginalis
Mycobacteroides abscessus
Mycoplasmopsis fermentans

Mycoplasmopsis primatum
Negativicoccus succinicivorans
Nevskia ramosa
Nosocomiicoccus ampullae
Odoribacter splanchnicus
Olegusella massiliensis
Oligella urethralis
Paenibacillus pabuli
Pantoea agglomerans
Parabacteroides distasonis
Parabacteroides merdae
Parachlamydia acanthamoebae
Paracoccus yeei
Parvimonas micra
Parvimonas parva
Pedobacter alluvionis
Pelomonas aquatica
Peptococcus niger
Peptoniphilus asaccharolyticus
Peptoniphilus gorbachii
Peptoniphilus grossensis
Peptoniphilus harei
Peptoniphilus koenoeneniae
Peptoniphilus lacrimalis
Peptoniphilus raoultii
Peptoniphilus timonensis
Peptostreptococcus anaerobius
Peptostreptococcus stomatis
Phocaeicola dorei
Photobacterium angustum
Porphyromonas asaccharolytica
Porphyromonas bennonis
Porphyromonas gingivalis
Porphyromonas somerae
Porphyromonas uenonis
Prevotella bivia
Prevotella brunnea
Prevotella corporis
Prevotella illustrans
Prevotella melaninogenica
Propionimicrobium lymphophilum
Pseudoclavibacter alba
Pseudoglutamicibacter cumminsii
Pseudomonas aeruginosa

Pseudomonas denitrificans
Pseudomonas mendocina
Pseudomonas monteilii
Pseudomonas oleovorans
Pseudomonas oryzihabitans
Pseudoramibacter alactolyticus
Pseudoroseomonas cervicalis
Rhizobium etli
Rhizorhabdus wittichii
Rhodanobacter terrae
Rothia kristinae
Rothia mucilaginosa
Ruminococcus flavefaciens
Ruminococcus torques
Salmonella enterica
Schaalia radingae
Sediminibacterium salmoneum
Segatella copri
Shewanella algae
Sneathia sanguinegens
Solobacterium moorei
Sphingobium yanoikuyae
Sphingomonas asaccharolytica
Staphylococcus aureus
Staphylococcus capitis
Staphylococcus caprae
Staphylococcus haemolyticus
Staphylococcus pettenkoferi
Staphylococcus saprophyticus
Staphylococcus simulans
Staphylococcus warneri
Staphylococcus xylosus
Stenotrophomonas maltophilia
Streptococcus agalactiae
Streptococcus anginosus
Streptococcus constellatus
Streptococcus gallolyticus
Streptococcus parasanguinis
Streptococcus sanguinis
Streptococcus thermophilus
Sutterella stercoricanis
Sutterella wadsworthensis
Treponema phagedenis
Ureaplasma urealyticum

Varibaculum cambriense
Variovorax paradoxus
Veillonella atypica
Veillonella montpellierensis
Veillonella parvula
Vibrio cholerae
Weeksella virosa
Winkia neuii
Xanthomonas campestris
Yersinia enterocolitica
