## Supplementary Table 11 for "HUGMi: Human Uro-Genital Microbiome database and hybrid classifier for improved species level annotation of 16S rRNA amplicon sequences"

**Supplementary Table 11: Comparison of taxonomic assignment with the actual stu**

### Comparison with taxa reported in study

#### Genus

| From dataset II | updated names | HUGMi assignment |
|---|---|---|
| Dialister | same | Dialister |
| Gardnerella | same | Gardnerella |
| Lactobacillus | same | Lactobacillus |
| Roseomonas | same | -- |
| Sneathia | same | Sneathia |
| Staphylococcus | same | Staphylococcus |
| Streptococcus | same | Streptococcus |

#### Species

| From dataset II | updated names | HUGMi assignment |
|---|---|---|
| Atopobium parvulum | Lancefieldella pa | -- |
| Gardnerella vaginalis | same | Gardnerella vaginalis |
| Lactobacillus crispat | same | Lactobacillus crispatus |
| Lactobacillus iners | same | Lactobacillus iners |
| Prevotella denticola | same | Prevotella denticola |
| Sneathia amnii | Sneathia vagina | Sneathia vaginalis |
| Streptococcus sangu | same | -- |

### Comparison with taxa by closed-reference OTU picking with Greeengenes

#### Genus

| From dataset II | updated names | HUGMi assignment |
|---|---|---|
| Acinetobacter | same | Acinetobacter |
| Actinobaculum | same | Actinobaculum |
| Actinomyces | same | Actinomyces |
| Aerococcus | same | Aerococcus |
| Alloscardovia | same | Alloscardovia |
| Anaerococcus | same | Anaerococcus |
| Anoxybacillus | same | -- |
| Arcanobacterium | same | Arcanobacterium |
| Atopobium | same | Atopobium |
| Bacillus | same | Bacillus |
| Bacteroides | same | Bacteroides |
| Bifidobacterium | same | Bifidobacterium |
| Bilophila | same | Bilophila |

| | | |
|---|---|---|
| Blautia | same | Blautia |
| Brevibacterium | same | Brevibacterium |
| Bulleidia | same | Bulleidia |
| Campylobacter | same | Campylobacter |
| Clostridium | same | Clostridium |
| Collinsella | same | Collinsella |
| Corynebacterium | same | Corynebacterium |
| Curtobacterium | same | -- |
| Dermabacter | same | Dermabacter |
| Dialister | same | Dialister |
| Dorea | same | Dorea |
| Enterococcus | same | Enterococcus |
| Facklamia | same | -- |
| Faecalibacterium | same | Faecalibacterium |
| Finegoldia | same | Finegoldia |
| Fusobacterium | same | Fusobacterium |
| Gallicola | same | Gallicola |
| Gardnerella | same | Gardnerella |
| Gemella | same | Gemella |
| Granulicatella | same | Granulicatella |
| Haemophilus | same | Haemophilus |
| Helcococcus | same | -- |
| Jonquetella | same | Jonquetella |
| Klebsiella | same | Klebsiella |
| Kocuria | same | -- |
| Lachnospira | same | Lachnospira |
| Lactobacillus | same | Lactobacillus |
| Lactococcus | same | Lactococcus |
| Megasphaera | same | Megasphaera |
| Micrococcus | same | Micrococcus |
| Mobiluncus | same | Mobiluncus |
| Mogibacterium | same | Mogibacterium |
| Moryella | same | Moryella |
| Mycoplasma | same | -- |
| Neisseria | same | Neisseria |
| Oligella | same | Oligella |
| Parvimonas | same | Parvimonas |
| Peptococcus | same | Peptococcus |
| Peptoniphilus | same | Peptoniphilus |
| Peptostreptococcus | same | Peptostreptococcus |
| Porphyromonas | same | Porphyromonas |
| Prevotella | same | Prevotella |
| Roseburia | same | Roseburia |
| Ruminococcus | same | Ruminococcus |

| | | |
|---|---|---|
| Shuttleworthia | same | -- |
| Sneathia | same | Sneathia |
| Sphingomonas | same | Sphingomonas |
| Staphylococcus | same | Staphylococcus |
| Streptococcus | same | Streptococcus |
| Succinivibrio | same | -- |
| Sutterella | same | Sutterella |
| Ureaplasma | same | Ureaplasma |
| Varibaculum | same | Varibaculum |
| Veillonella | same | Veillonella |
| | | Acidaminococcus |
| | | Acidovorax |
| | | Actinotignum |
| | | Aedoeadaptatus |
| | | Aeromonas |
| | | Agathobacter |
| | | Akkermansia |
| | | Alistipes |
| | | Alloprevotella |
| | | Altererythrobacter |
| | | Anaeroglobus |
| | | Barnesiella |
| | | Capnocytophaga |
| | | Catonella |
| | | Chlamydia |
| | | Citrobacter |
| | | Cloacibacterium |
| | | Clostridioides |
| | | Coprococcus |
| | | Dechloromonas |
| | | Desulfovibrio |
| | | Eggerthella |
| | | Eggerthia |
| | | Enterobacter |
| | | Eremococcus |
| | | Escherichia |
| | | Eubacterium |
| | | Ezakiella |
| | | Fannyhessea |
| | | Fastidiosipila |
| | | Fenollaria |
| | | Filifactor |
| | | Flavonifractor |
| | | Gemelliphila |

Gordonibacter
Hallella
Howardella
Hoylesella
Jeotgalicoccus
Khoudiadiopia
Kingella
Lacticaseibacillus
Lawsonella
Leclercia
Levilactobacillus
Leyella
Ligilactobacillus
Limosilactobacillus
Mageeibacillus
Massilia
Mediterraneibacter
Metamycoplasma
Methylobacterium
Methylorubrum
Moraxella
Murdochiella
Mycobacteroides
Mycoplasmopsis
Negativicoccus
Nosocomiicoccus
Olegusella
Olsenella
Paenibacillus
Pantoea
Parabacteroides
Paracoccus
Pediococcus
Phocaeicola
Photobacterium
Propionimicrobium
Pseudoglutamicibacter
Pseudomonas
Rhodanobacter
Rothia
Salmonella
Schaalia
Sediminibacterium
Segatella

| | | |
|---|---|---|
| | | Shewanella |
| | | Solobacterium |
| | | Thomasclavelia |
| | | Treponema |
| | | Trueperella |
| | | Turicibacter |
| | | Virgibacillus |
| | | Xanthomonas |
| | | Yersinia |

| **Species** | | |
|---|---|---|
| **From dataset II** | **updated names** | **HUGMi assignment** |
| Acinetobacter johnsc | same | -- |
| Acinetobacter lwoffii | same | Acinetobacter lwoffii |
| Anoxybacillus kestar | same | -- |
| Bacteroides ovatus | same | Bacteroides ovatus |
| Bacteroides uniformi | same | -- |
| Bifidobacterium adol | same | Bifidobacterium adolescentis |
| Bifidobacterium long | same | Bifidobacterium longum |
| Blautia obeum | same | Blautia obeum |
| Campylobacter ureo | same | Campylobacter ureolyticus |
| Clostridium perfringe | same | Clostridium perfringens |
| Collinsella aerofacie | same | Collinsella aerofaciens |
| Dorea formicigenera | same | Dorea formicigenerans |
| Faecalibacterium pra | same | Faecalibacterium prausnitzii |
| Atopobium vaginae | Fannyhessea va | Fannyhessea vaginae |
| Actinomyces europa | Gleimia europae | -- |
| Haemophilus parainf | same | Haemophilus parainfluenzae |
| Jonquetella anthropi | same | Jonquetella anthropi |
| Kocuria rhizophila | same | -- |
| Lactobacillus iners | same | Lactobacillus iners |
| Prevotella stercorea | Leyella stercorea | Leyella stercorea |
| Lactobacillus ruminis | Ligilactobacillus | -- |
| Lactobacillus coleoh | Limosilactobacil | Limosilactobacillus coleohominis |
| Lactobacillus reuteri | Limosilactobacil | Limosilactobacillus reuteri |
| Lactobacillus vagina | Limosilactobacil | -- |
| Moryella indoligenes | same | Moryella indoligenes |
| Neisseria subflava | same | -- |
| Peptostreptococcus | same | Peptostreptococcus anaerobius |
| Prevotella melaninog | same | Prevotella melaninogenica |
| Prevotella copri | Segatella copri | Segatella copri |
| Bulleidia moorei | Solobacterium m | Solobacterium moorei |
| Staphylococcus aure | same | Staphylococcus aureus |
| Staphylococcus epid | same | Staphylococcus epidermidis |

| | | |
|---|---|---|
| Staphylococcus hae | same | Staphylococcus haemolyticus |
| Streptococcus agala | same | Streptococcus agalactiae |
| Streptococcus angin | same | Streptococcus anginosus |
| Streptococcus infant | same | -- |
| Veillonella dispar | same | -- |
| Veillonella parvula | same | Veillonella parvula |
| | | Acidaminococcus intestini |
| | | Acidaminococcus massiliensis |
| | | Acidovorax delafieldii |
| | | Acinetobacter baumannii |
| | | Actinobaculum massiliense |
| | | Actinomyces oris |
| | | Actinomyces provencensis |
| | | Actinotignum schaalii |
| | | Actinotignum urinale |
| | | Aedoeadaptatus coxii |
| | | Aerococcus sanguinicola |
| | | Aeromonas caviae |
| | | Aeromonas enteropelogenes |
| | | Agathobacter rectalis |
| | | Akkermansia muciniphila |
| | | Alistipes finegoldii |
| | | Alistipes shahii |
| | | Alloprevotella tannerae |
| | | Alloscardovia omnicolens |
| | | Altererythrobacter epoxidivorans |
| | | Anaerococcus hydrogenalis |
| | | Anaerococcus jeddahensis |
| | | Anaerococcus lactolyticus |
| | | Anaerococcus prevotii |
| | | Anaerococcus provencensis |
| | | Anaerococcus tetradius |
| | | Anaerococcus vaginimassiliensis |
| | | Anaeroglobus geminatus |
| | | Arcanobacterium haemolyticum |
| | | Arcanobacterium ihumii |
| | | Atopobium minutum |
| | | Barnesiella viscericola |
| | | Bifidobacterium animalis |
| | | Bifidobacterium bifidum |
| | | Bifidobacterium breve |
| | | Bifidobacterium dentium |
| | | Bilophila wadsworthia |
| | | Brevibacterium ravenspurgense |

Bulleidia extructa
Campylobacter gracilis
Campylobacter hominis
Capnocytophaga ochracea
Catonella morbi
Chlamydia trachomatis
Citrobacter werkmanii
Cloacibacterium normanense
Clostridioides difficile
Clostridium butyricum
Coprococcus eutactus
Corynebacterium aurimucosum
Corynebacterium genitalium
Corynebacterium glucuronolyticum
Corynebacterium lipophilum
Corynebacterium otitidis
Corynebacterium pseudokroppenstedtii
Corynebacterium riegelii
Corynebacterium striatum
Corynebacterium urealyticum
Dechloromonas agitata
Dermabacter hominis
Dermabacter vaginalis
Desulfovibrio desulfuricans
Desulfovibrio fairfieldensis
Dialister invisus
Dialister micraerophilus
Dialister pneumosintes
Dialister propionicifaciens
Dorea longicatena
Eggerthella lenta
Eggerthia catenaformis
Enterobacter bugandensis
Enterobacter cloacae
Enterococcus mundtii
Enterococcus pseudoavium
Eremococcus coleocola
Escherichia coli
Escherichia fergusonii
Eubacterium nodatum
Eubacterium saphenum
Eubacterium yurii
Ezakiella coagulans
Ezakiella massiliensis

Fastidiosipila sanguinis
Fenollaria massiliensis
Filifactor alocis
Finegoldia magna
Flavonifractor plautii
Fusobacterium animalis
Fusobacterium gonidiaformans
Fusobacterium nucleatum
Fusobacterium periodonticum
Gallicola barnesae
Gemella haemolysans
Gemelliphila asaccharolytica
Gordonibacter pamelaeae
Granulicatella elegans
Hallella bergensis
Hallella colorans
Howardella ureilytica
Hoylesella marshii
Hoylesella nanceiensis
Hoylesella timonensis
Jeotgalicoccus psychrophilus
Khoudiadiopia massiliensis
Kingella kingae
Klebsiella aerogenes
Klebsiella pneumoniae
Lachnospira eligens
Lacticaseibacillus rhamnosus
Lactobacillus delbrueckii
Lactobacillus gasseri
Lactobacillus jensenii
Lactobacillus johnsonii
Lactococcus lactis
Lawsonella clevelandensis
Leclercia adecarboxylata
Levilactobacillus brevis
Ligilactobacillus salivarius
Limosilactobacillus mucosae
Mageeibacillus indolicus
Massilia timonae
Mediterraneibacter gnavus
Megasphaera hutchinsoni
Metamycoplasma hominis
Metamycoplasma salivarium
Methylobacterium radiotolerans

Methylorubrum zatmanii
Micrococcus luteus
Mobiluncus curtisii
Mobiluncus mulieris
Mogibacterium timidum
Moraxella catarrhalis
Moraxella osloensis
Murdochiella asaccharolytica
Murdochiella vaginalis
Mycobacteroides abscessus
Mycoplasmopsis primatum
Negativicoccus succinicivorans
Neisseria meningitidis
Nosocomiicoccus ampullae
Olegusella massiliensis
Oligella urethralis
Olsenella urininfantis
Pantoea agglomerans
Parabacteroides merdae
Paracoccus yeei
Parvimonas micra
Parvimonas parva
Pediococcus acidilactici
Peptococcus niger
Peptoniphilus duerdenii
Peptoniphilus gorbachii
Peptoniphilus koenoeneniae
Peptoniphilus lacrimalis
Peptoniphilus timonensis
Peptostreptococcus stomatis
Phocaeicola dorei
Phocaeicola vulgatus
Photobacterium angustum
Porphyromonas asaccharolytica
Porphyromonas bennonis
Porphyromonas endodontalis
Porphyromonas somerae
Porphyromonas uenonis
Prevotella bivia
Prevotella brunnea
Prevotella corporis
Prevotella histicola
Prevotella ihumii
Prevotella melaninogenica

Prevotella nigrescens
Prevotella timonensis
Prevotella veroralis
Propionimicrobium lymphophilum
Pseudoglutamicibacter cumminsii
Pseudomonas aeruginosa
Pseudomonas alcaligenes
Pseudomonas oryzihabitans
Rhodanobacter terrae
Roseburia inulinivorans
Rothia mucilaginosa
Ruminococcus flavefaciens
Salmonella enterica
Schaalia radingae
Sediminibacterium salmoneum
Segatella oris
Shewanella putrefaciens
Sneathia sanguinegens
Sphingomonas asaccharolytica
Sphingomonas paucimobilis
Staphylococcus caprae
Staphylococcus warneri
Streptococcus dysgalactiae
Streptococcus mutans
Streptococcus oralis
Streptococcus salivarius
Streptococcus thermophilus
Sutterella stercoricanis
Sutterella wadsworthensis
Thomasclavelia ramosa
Treponema phagedenis
Trueperella bernardiae
Turicibacter sanguinis
Ureaplasma urealyticum
Varibaculum cambriense
Veillonella atypica
Virgibacillus proomii
Xanthomonas campestris
Yersinia enterocolitica

udy (dataset II)
