## Supplementary Table 12 for "HUGMi: Human Uro-Genital Microbiome database and hybrid classifier for improved species level annotation of 16S rRNA amplicon sequences"

**Supplementary Table 12: Differentially significant bacterial species identified by HUGMi**

**Significant taxa for dataset I**

| Species-level taxa | base Mean | log2Fold Change | lfcSE | stat | pvalue | padj | Group |
|---|---|---|---|---|---|---|---|
| Veillonella parvula | 123.17 | -7.43015 | 1.161285 | -6.3982 | 1.57E-10 | 8.80E-09 | Bladder cancer |
| Fusobacterium nucleatum | 622.04 | -6.95699 | 1.282981 | -5.4225 | 5.88E-08 | 1.57E-06 | Bladder cancer |
| Jonquetella anthropi | 50.719 | -6.1406 | 1.136455 | -5.4033 | 6.54E-08 | 1.57E-06 | Bladder cancer |
| Streptococcus agalactiae | 25.651 | -5.28674 | 1.141113 | -4.633 | 3.60E-06 | 6.34E-05 | Bladder cancer |
| Campylobacter hominis | 36.13 | -5.11138 | 1.105493 | -4.6236 | 3.77E-06 | 6.34E-05 | Bladder cancer |
| Yersinia enterocolitica | 8.5509 | -3.48854 | 0.912552 | -3.8228 | 0.000132 | 0.00123 | Bladder cancer |
| Escherichia coli | 172.86 | -4.1238 | 1.136252 | -3.6293 | 0.000284 | 0.00239 | Bladder cancer |
| Sphingomonas asaccharolytica | 18.885 | -3.15741 | 1.01234 | -3.1189 | 0.001815 | 0.01215 | Bladder cancer |
| Catonella morbi | 38.563 | -3.16509 | 1.027255 | -3.0811 | 0.002062 | 0.01242 | Bladder cancer |
| Alistipes finegoldii | 7.0264 | -2.47714 | 0.83986 | -2.9495 | 0.003183 | 0.01783 | Bladder cancer |
| Leclercia adecarboxylata | 290.62 | -3.32996 | 1.175186 | -2.8336 | 0.004603 | 0.02417 | Bladder cancer |
| Murdochiella vaginalis | 285.38 | -3.4113 | 1.256156 | -2.7157 | 0.006614 | 0.03268 | Bladder cancer |
| Escherichia fergusonii | 63.377 | -3.08936 | 1.180575 | -2.6168 | 0.008875 | 0.0403 | Bladder cancer |
| Veillonella atypica | 17816 | 8.331265 | 1.424003 | 5.85059 | 4.90E-09 | 2.06E-07 | Healthy |
| Corynebacterium provencense | 31.537 | 5.227362 | 0.982942 | 5.31808 | 1.05E-07 | 2.20E-06 | Healthy |
| Enterococcus mundtii | 63.963 | 4.401657 | 1.018904 | 4.31999 | 1.56E-05 | 0.0002 | Healthy |
| Paracoccus yeei | 14.137 | 3.455454 | 0.851862 | 4.05636 | 4.98E-05 | 0.00056 | Healthy |
| Gemella haemolysans | 19.966 | 2.938842 | 1.014818 | 2.89593 | 0.00378 | 0.02049 | Healthy |
| Parvimonas parva | 171.54 | 3.071256 | 1.210142 | 2.53793 | 0.011151 | 0.0479 | Healthy |

**Significant taxa for dataset II**

| Species-level taxa | base Mean | log2Fold Change | lfcSE | stat | pvalue | padj | Group |
|---|---|---|---|---|---|---|---|
| Gemella haemolysans | 10.613 | -3.88762 | 0.830036 | -4.6837 | 2.82E-06 | 0.00015 | Cervical cancer |
| Sneathia sanguinegens | 22.55 | -4.24792 | 0.935191 | -4.5423 | 5.56E-06 | 0.0002 | Cervical cancer |
| Peptostreptococcus stomatis | 5.938 | -2.71176 | 0.791985 | -3.424 | 0.000617 | 0.00617 | Cervical cancer |
| Pseudomonas aeruginosa | 97.654 | -3.29596 | 1.047453 | -3.1466 | 0.001652 | 0.01069 | Cervical cancer |
| Parvimonas parva | 10.938 | -2.63257 | 0.871597 | -3.0204 | 0.002524 | 0.01341 | Cervical cancer |
| Staphylococcus warneri | 15.047 | -2.00366 | 0.804457 | -2.4907 | 0.012749 | 0.0425 | Cervical cancer |
| Fusobacterium nucleatum | 20.991 | 4.591902 | 1.073481 | 4.27758 | 1.89E-05 | 0.00035 | Healthy |
| Dialister propionicifaciens | 73.244 | 4.185843 | 1.05711 | 3.95971 | 7.50E-05 | 0.00118 | Healthy |
| Aedoeadaptatus coxii | 10.146 | 3.389942 | 0.880707 | 3.84912 | 0.000119 | 0.00163 | Healthy |
| Streptococcus anginosus | 33.3 | 3.900098 | 1.031943 | 3.77937 | 0.000157 | 0.00192 | Healthy |
| Catonella morbi | 9.1189 | 3.615745 | 0.994595 | 3.6354 | 0.000278 | 0.00305 | Healthy |

| | | | | | | | |
|---|---|---|---|---|---|---|---|
| Campylobacter ureolyticus | 15.621 | 3.063842 | 0.940144 | 3.25891 | 0.001118 | 0.0082 | Healthy |
| Peptoniphilus lacrimalis | 4.5633 | 2.551362 | 0.839939 | 3.03756 | 0.002385 | 0.01341 | Healthy |
| Peptoniphilus koenoeneniae | 4.7871 | 2.627401 | 0.867697 | 3.02802 | 0.002462 | 0.01341 | Healthy |
| Lactobacillus jensenii | 416.7 | 3.778931 | 1.32813 | 2.8453 | 0.004437 | 0.01861 | Healthy |
| Acinetobacter baumannii | 5.3098 | 2.504585 | 0.883122 | 2.83606 | 0.004567 | 0.01861 | Healthy |
| Prevotella bivia | 155.8 | 2.959546 | 1.155072 | 2.56222 | 0.010401 | 0.03691 | Healthy |
| Fenollaria massiliensis | 58.98 | 2.230831 | 1.061859 | 2.10087 | 0.035652 | 0.09565 | Healthy |
