## Supplementary Figures 1, 2 for "HUGMi: Human Uro-Genital Microbiome database and hybrid classifier for improved species level annotation of 16S rRNA amplicon sequences"

| | | Expected values | | |
|---|---|---|---|---|
| | | No classification | Under classification | Genus/Species level classification |
| Real values | Bacterial sequence | **FN** | **FN** | **TP** |
| | Random sequence | **TN** | **FP** | **FP** |

**Figure 1:** Confusion matrix depicting the TP (True Positive), TN (True Negative), FP (False Positive) and FN (False Negative) values of taxonomic classification.

$$Precision = \frac{TP}{TP + FP} \quad (1)$$

$$Recall = \frac{TP}{TP + FN} \quad (2)$$

$$F1 = 2 \times \frac{Precision \times Recall}{Precision + Recall} \quad (3)$$

**Figure 2:** Equations used for the calculation of Precision, Recall and F1 score.
