## Supplementary Table 1 for "HUGMi: Human Uro-Genital Microbiome database and hybrid classifier for improved species level annotation of 16S rRNA amplicon sequences"

**Supplementary Table 1: Database information**

| Database name | No. of initial taxa | No. of taxa after filtering in step (I.) |
|---|---|---|
| EzBioCloud | 64660 | 15403 |
| GTDB | 584382 | 320880 |
| NCBI | 22061 | 21079 |
| SILVA | 510984 | 127738 |
| RDP | 3356808 | 191086 |
| GG2 | 331269 | 134803 |
